# The lysosomal membrane protein LAMP2B mediates microlipophagy to target obesity-related disorders

**DOI:** 10.1101/2024.09.18.613587

**Authors:** Ryohei Sakai, Shu Aizawa, Hyeon-Cheol Lee-Okada, Katsunori Hase, Hiromi Fujita, Hisae Kikuchi, Yukiko U. Inoue, Takayoshi Inoue, Chihana Kabuta, Takehiko Yokomizo, Tadafumi Hashimoto, Keiji Wada, Tatsuo Mano, Ikuko Koyama-Honda, Tomohiro Kabuta

## Abstract

Lifestyle diseases, such as obesity, diabetes, and metabolic syndrome, are leading health problems, most of which are related to abnormal lipid metabolism. Lysosomes can degrade lipid droplets (LDs) via microautophagy. Here, we report the molecular mechanism and pathophysiological roles of microlipophagy, regulated by the lysosomal membrane protein LAMP2B. Our study revealed that LAMP2B interacts with phosphatidic acid, facilitating lysosomal-LD interactions and enhancing lipid hydrolysis via microlipophagy depending on endosomal sorting complexes required for transport. Correlative light and electron microscopy demonstrated direct LDs uptake into lysosomes at contact sites. Moreover, LAMP2B overexpression in mice prevents high-fat diet-induced obesity, insulin resistance, and adipose tissue inflammation; liver lipidomics analysis suggested enhanced triacylglycerol hydrolysis. Overall, the findings of this study elucidated the mechanism of microlipophagy, which could be promising for the treatment of obesity and related disorders.

## INTRODUCTION

Recently, there has been a steady increase in the number of patients with lifestyle diseases, such as obesity and diabetes mellitus.^1^ Several diseases related to abnormal lipid metabolism, including chronic kidney disease, fatty liver, heart failure, obesity, neurodegeneration, and cancer, are at least partly caused by the intracellular accumulation of triacylglycerols (TAGs) and other neutral lipids in lipid droplets (LDs).^2–4^ Therefore, understanding the lipid degradation system is essential in elucidating lipid metabolism disorders^2–4^. The hydrolysis of TAGs and cholesteryl esters in LDs is thought to involve two central catabolic pathways: cytosolic lipolysis and macroautophagy.^5,6^

There are at least three classes of autophagy: macroautophagy, chaperone-mediated autophagy (CMA), and microautophagy.^7,8^ Macroautophagy is characterized by a lipid double-membrane enclosing cytoplasmic components and/or organelles, which eventually fuses with lysosomes where its contents are degraded. Degradation of LDs via macroautophagy is termed as macrolipophagy, which involves transporting intracellular LDs sequestered in double-membraned vesicles (autophagosomes) to lysosomes for degradation.^9^ In CMA, proteins containing the KFERQ (-like) motif are directly transported into lysosomes via HSC70 and a translocation complex with LAMP2A on the lysosomal membrane. Microautophagy involves the direct uptake of cytosolic compartment or organelles through invagination of the lysosomal membrane. Studies have revealed that CMA selectively degrades the LD-associated proteins perilipin 2 (PLIN2) and perilipin 3 (PLIN3) in lysosomes, exposing the lipid core to adipose triglyceride lipase (ATGL) activity. Additionally, the elimination of PLINs allows proteins involved in autophagosome formation (ATGs) to induce macrolipophagy.^10^

Recently, microlipophagy has been reported in mammalian hepatocytes; hepatocyte lysosomes and LDs undergo interactions during which proteins and lipids can be transferred from LDs directly into lysosomes.^11^ However, molecular tethers or regulatory proteins that promote the bridging of the lysosome and LDs have not been identified, and the underlying molecular mechanisms are unknown.

Recent studies have revealed that organelle-organelle interactions are mediated by protein-phospholipid or protein-protein interactions. The protein OPA1 and the phospholipid cardiolipin function together in mitochondrial inner membrane fusion,^12^ and LDs bind to several organelles via protein-protein interactions.^13^ Therefore, we hypothesized that protein-protein or protein-phospholipid interactions are involved in the interaction between lysosomes and LDs during microlipophagy.

One of the lysosomal membrane proteins that interacts with phospholipids or LD proteins is LAMP2B. LAMP2 is a major lysosomal membrane protein with three variants produced from alternative splicing: LAMP2A, LAMP2B, and LAMP2C.^14^ These LAMPs possess a cytosolic sequence of approximately 11 amino acids at the C-terminus. The cytosolic sequence of LAMP2A interacts with HSC70 during CMA^15^ and that of LAMP2C can interact with substrate nucleic acids during RNautophagy and DNautophagy, unconventional types of autophagy in which nucleic acids are directly taken up by lysosomes.^16–19^ The cytosolic sequence of LAMP2B interacted with purified nucleic acids,^18^ but did not interact with RNA in mouse brain lysate, which was solubilized with the detergent Triton-X 100.^17^ Because Triton-X 100 solubilizes membrane lipids, we speculated that the cytosolic sequence of LAMP2B interacts with lipid molecules and might be involved in lipid hydrolysis. Therefore, in this study, we examined the role of LAMP2B in lipid hydrolysis, focusing on microlipophagy.

## RESULTS

### Cytosolic region of LAMP2B interacts with phosphatidic acid

LAMP2A, B, and C possess distinct cytosolic sequences of approximately 11 amino acids at the C-terminus (Figure 1A). We examined the inhibitory effects of different types of lipids on the interaction between purified RNA and LAMP2B peptide (Figure 1A). The results of a pull-down assay showed that the addition of phosphatidic acid (PA) inhibited the interaction between the LAMP2B peptide and purified RNA (Figure 1B), whereas the addition of phosphatidylcholine or cholesterol did not (Figure 1B); this suggests that the cytosolic sequence of LAMP2B possesses affinity for PA. The interaction between LAMP2B and PA was confirmed with pull-down and lipid array assays (Figure 1C, D). Additionally, the cytoplasmic sequence of LAMP2C interacted with PA, whereas that of LAMP2A did not (Figure 1D).

**Figure 1.**
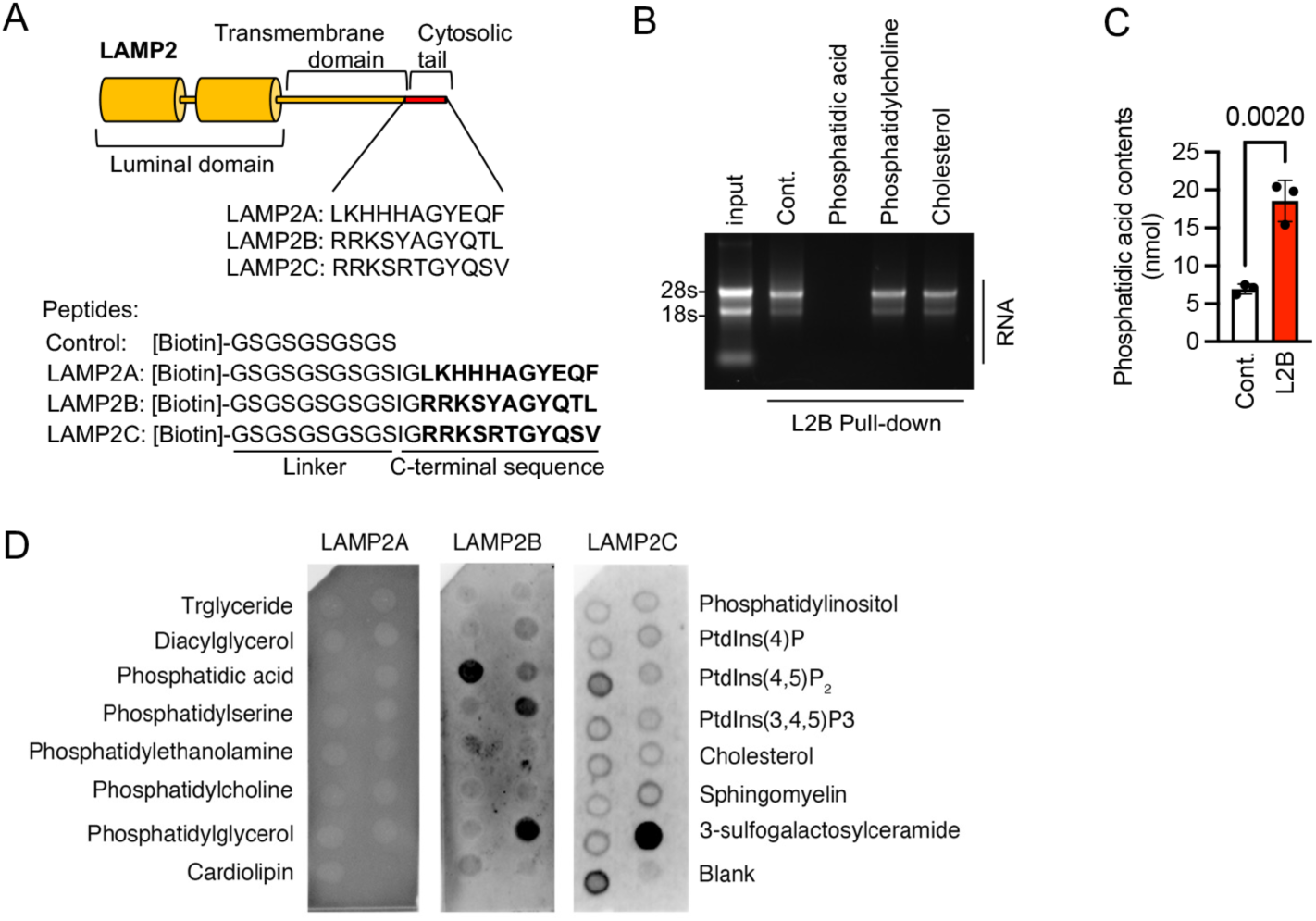
Cytosolic region of LAMP2B interacts with phosphatidic acid. (A) Schematic of LAMP2s and their cytosolic sequences (upper panel) and biotin-tagged peptides for LAMP2s cytosolic tail (lower panel). (B) Competitive inhibition of RNA binding of LAMP2B (L2B) cytosolic tails by lipid molecules. (C) Interaction of LAMP2B cytosolic tail and PA by pull-down assay. Data are presented as mean ± SD (n = 3). P-values are obtained from unpaired *t* test. (D) Membrane lipid binding assay of LAMP2 cytosolic tails.

### LAMP2B decreases cellular LD levels via lysosomal acid lipase, and not cytosolic lipase

Recent studies have revealed that PA is involved in the fusion or transport of LDs via PA-protein interactions. For instance, the CIDEA protein promotes LD fusion by binding to PA.^20^ Furthermore, PA on LD interacts with kinesin-1, which transports LD toward the ER in hepatocytes.^21^ Therefore, we hypothesized that LAMP2B in lysosomes may interact with PA in LDs and mediate lysosomal hydrolysis of lipid molecules (TAGs and cholesteryl esters).

To confirm this, we examined the effect of siRNA-mediated knockdown (KD) of LAMP2B, LAMP2A, and LAMP2C on the hydrolysis of lipid molecules using previously described methods.^22–24^ HeLa cells, which have been used to study LD degradation,^25^ were used for this experiment. The TAGs in control and KD cells were labeled with oleic acid containing [^3^H] oleic acid, after which the efflux of [^3^H] oleic acid into the medium was monitored for 3 h. Triacsin C was included in all incubations during the efflux phase to prevent the reutilization of fatty acids released from hydrolyzed TAG (Figure 2A). The results showed that KD of LAMP2B and LAMP2C decreased the extracellular radioactivity derived from the labeled free fatty acids, indicating decreased lipid hydrolysis (Figure 2B, 2C, S1A, S1B, S1D, and S1E). Moreover, Oil Red O staining (Figure 2A) showed that KD of LAMP2B and LAMP2C induced significant LD accumulation in the cells, whereas KD of LAMP2A did not (Figure 2D, 2E and S1A-S1E). However, KD of LAMP2A decreased lipid hydrolysis (Figure 2B, S1A, S1C); this is consistent with previous findings^10^ that have shown that LAMP2A promotes the degradation of LDs via the cytosolic lipase ATGL through CMA-mediated degradation of the LD-associated PLIN2 and PLIN3.^10^ Furthermore, the effect of overexpression of LAMP2 proteins on lipid hydrolysis was examined using [^3^H] oleic acid (Figure 2F). LAMP2B overexpression markedly increased lipid hydrolysis, whereas LAMP2A overexpression did not (Figure 2G, S1F). Similarly, LAMP2C overexpression increased lipid hydrolysis, but to a lesser degree than LAMP2B overexpression (Figure 2G, S1F). The effect of overexpressing LAMP2 proteins on LD accumulation was examined using Oil Red O staining. The cells were lipogenically stimulated with 200 µM oleic acid (Figure 2F), and the results showed that LAMP2B overexpression suppressed oleic acid-induced increase in cellular levels of LD (Figure 2H). Similarly, LAMP2C overexpression decreased LD levels, but to a lesser degree than LAMP2B overexpression (Figure 2H). Collectively, LAMP2B affected lipid hydrolysis and LD accumulation in both KD and overexpression experiments, indicating that LAMP2B plays an important role in decreasing cellular LD levels.

**Figure 2.**
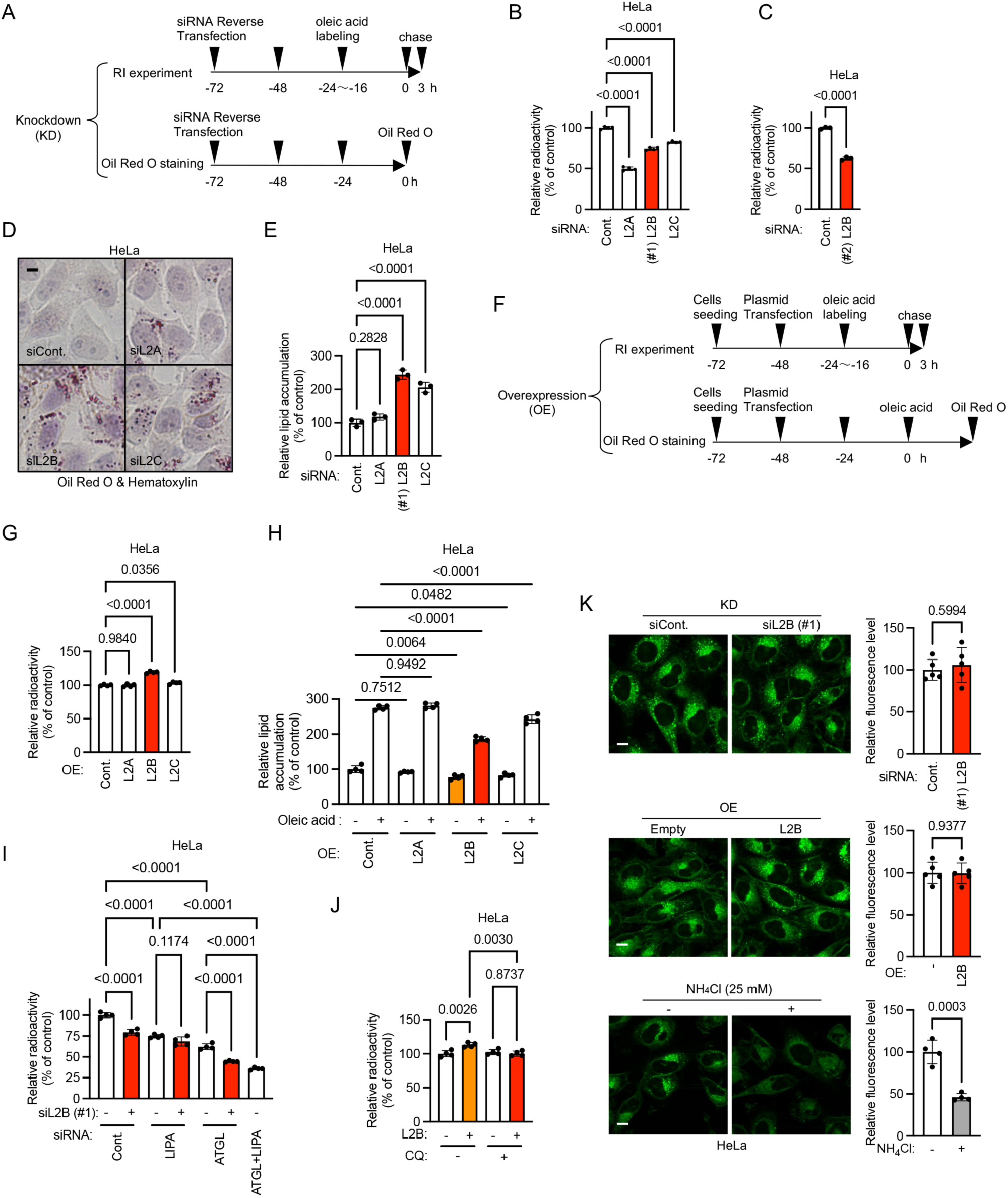
LAMP2B decreases cellular LD levels depending on lysosomal acid lipase, not on cytosolic lipase. (A) Schematics of the experimental procedures for lipid hydrolysis measurement, Oil Red O staining, and siRNA-KD. (B and C) Lipid hydrolysis level in LAMP2A, LAMP2B, LAMP2C-KD HeLa cells. Data are presented as mean ± SD (n = 3–4). (D) HeLa cells were transfected with *LAMP2A*, *LAMP2B*, or *LAMP2C* or *EGFP*-#2 siRNA, and neutral lipid deposit was evaluated using Oil Red O and hematoxylin stains. Scale bars: 10 µm. (E) Oil red O stain retention by *LAMP2A*-, *LAMP2B*-, or *LAMP2C*-specific siRNA cells was determined using a spectrophotometer. Data are presented as mean ± SD. (n = 3). (F) Schematics of the experimental procedures for RI experiment, Oil Red O staining of cells overexpressing the LAMP2 proteins. (G) Lipid hydrolysis level of LAMP2A, LAMP2B, LAMP2C-overexpressing HeLa cells. Data are presented as mean ± SD. (n = 4). (H) Oil red O-based quantification of LD accumulation in HeLa cells overexpressed with indicated LAMP2s. Data are presented as mean ± SD (n = 4). (I) Hela cells were transfected with *LAMP2B* and *LIPA* or *ATGL*-siRNA and lipid hydrolysis level was measured. Data are presented as mean ± SD (n = 4). (J) Lipid hydrolysis level of LAMP2B-overexpressing cells treated with or without CQ. Data are presented as mean ± SD (n = 4). (K) Acidity of lysosomes in LAMP2B-KD or -overexpressing HeLa cells. The control group was treated with or without NH_4_Cl. Data are presented as mean ± SD (n = 4). Scale bars: 10 µm. P-values are from unpaired t test (C and K), Dunnett’s multiple comparisons test (B, F and H) or Tukey’s multiple comparisons test (D, H, I, and J).

To investigate whether that LAMP2B-mediated lipid hydrolysis differs from cytosolic lipolysis, we examined the effect of double KD of LAMP2B and either lysosomal acid lipase (LIPA) or ATGL on lipid hydrolysis. The KD of LAMP2B, LIPA, or ATGL alone decreased lipid hydrolysis. The combined KD of ATGL and LIPA further decreased lipid hydrolysis compared to KD of ATGL or LIPA alone (Figure 2I and S1G), indicating the involvement of both lysosomal and cytosolic lipase pathways. Notably, the combined KD of LAMP2B and ATGL, but not LAMP2B and LIPA, significantly enhanced lipid hydrolysis compared to KD of ATGL alone or LIPA alone, respectively (Figure 2I and S1G), implying that LAMP2B-mediated lipid hydrolysis is lysosomal. To further confirm this result, the cells were treated with chloroquine (CQ), a lysosomal hydrolase inhibitor. LAMP2B overexpression did not increase lipid hydrolysis in cells treated with CQ (Figure 2J), indicating that LAMP2B indeed mediates lysosomal lipid hydrolysis. Additionally, the effect of LAMP2B on the pH of lysosomes was examined, and we found that neither overexpression nor KD of LAMP2B affected the pH of the lysosomes (Figure 2K). These results indicate that LAMP2B-mediated lipid hydrolysis is lysosomal (LIPA-dependent) and differs from cytosolic lipolysis.

### LAMP2B mediates microlipophagy

A recent study reported that microlipophagy was detected in primary rat hepatocytes and AML12 mouse hepatocytes,^11^ which possess several characteristics of human primary hepatocytes.^26^ Therefore, we investigated whether LAMP2B is involved in microlipophagy in AML12 cells. Our results showed that LAMP2B KD decreased lipid hydrolysis in AML12 cells (Figure 3A, S1H). Additionally, LAMP2B KD did not decrease lipid hydrolysis in cells treated with CQ (Figure 3B), indicating that LAMP2B mediates lysosomal lipid hydrolysis in AML12 cells, similar to HeLa cells (Figure 2). Next, we performed a colocalization experiment to determine whether LAMP2B is involved in the interaction between lysosomes and LDs, using a mCherry-GFP-Plin2 reporter that exhibits differential fluorescence depending on the pH of its surrounding environment (Figure 3C, S2A). Generally, the Plin2 reporter localizes to the surface of LDs and leaves behind small mCherry^+^-only puncta upon uptake by acidic lysosomes.^11,27^ In addition, all Alexa647-Dextran positive signals, indicative of lysosomes, were confirmed to co-localize with LAMP2B-GFP-positive signals (Figure S2B). Colocalization analysis revealed that LAMP2B KD did not affect the fraction of GFP (PLIN2 localized to LDs) overlapping dextran (lysosome), but decreased the fraction of mCherry (PLIN2 localized to LDs or lysosomes) overlapping dextran (Figure 3D and 3E), indicating decreased levels of PLIN2 in lysosomes in LAMP2B KD cells. Additionally, the colocalization of mCherry and GFP increased in LAMP2B KD cells (Figure 3D and 3F), indicating LD accumulation in LAMP2B KD cells.

**Figure 3.**
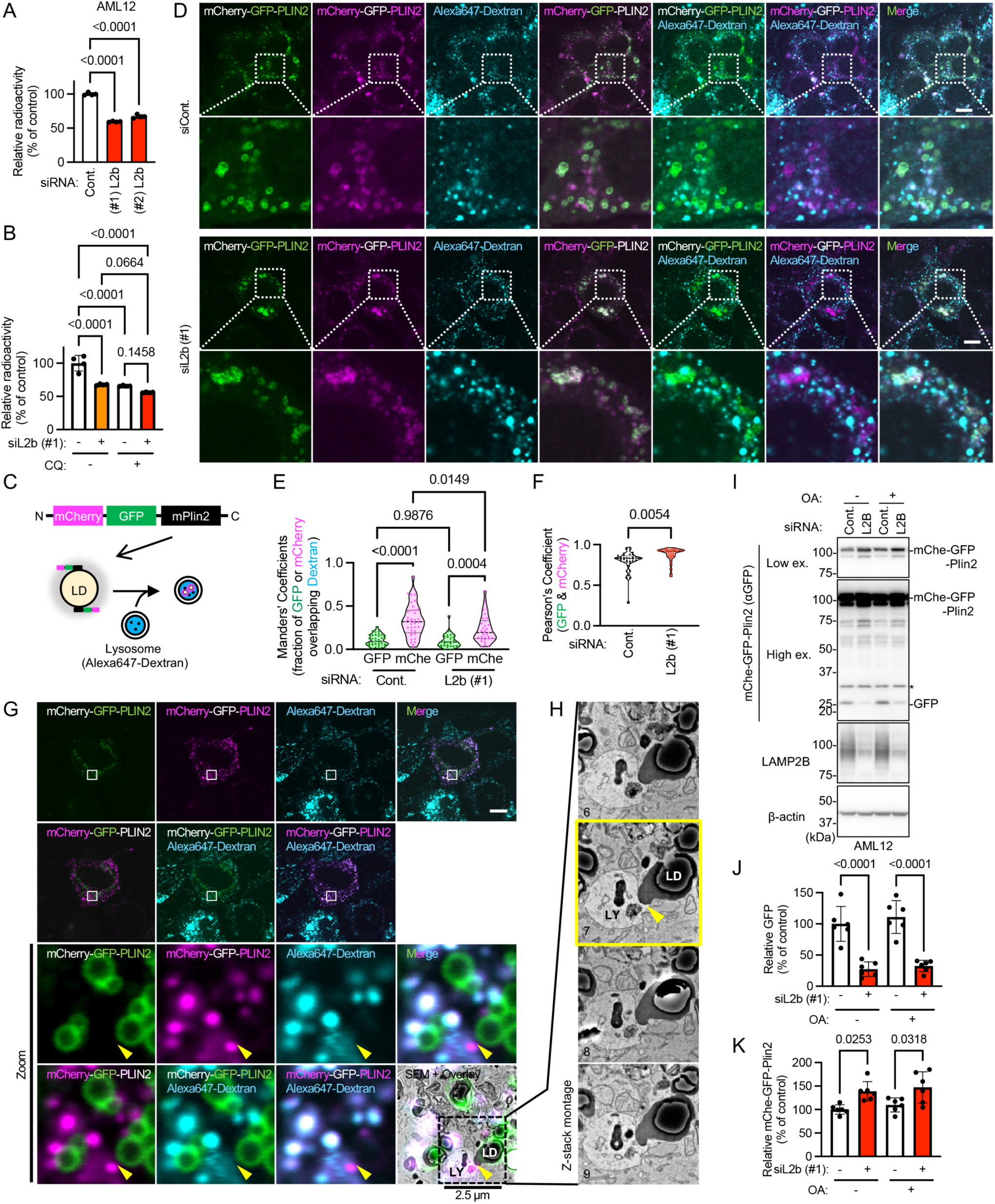
LAMP2B mediates microlipophagy. (A) Lipid hydrolysis level of Lamp2b-KD AML12 cells. Data are presented as mean ± SD (n = 4). (B) Lipid hydrolysis level of LAMP2B-KD cells treated with or without CQ. Data are presented as mean ± SD (n = 4) (C) Schematic of analysis of microlipophagy using the tandem-fluorescent perilipin reporter. N-terminal fusions of mCherry and EGFP to PLIN2 label the surface of cytoplasmic LD. Upon uptake of LD into an acidic organelle, such as a lysosome, the EGFP signal is selectively quenched, resulting in punctae only exhibiting mCherry fluorescence. Alexa647-dextran labeling the lumen of lysosome. Levels of colocalization of mCherry and Alexa647 indicate levels of LD inside lysosome. EGFP and Alexa647 exhibited minimal colocalization and were utilized as negative controls. Levels of colocalization of mCherry and EGFP indicate levels of cytoplasmic LD. (D) Cellular localization of mCherry-GFP-PLIN2 and Alexa647-Dextran in AML12 cells. Box framed regions are enlarged. Scale bars: 10 µm. (E) Manders’ correlation coefficient values quantify the levels of GFP or mCherry–Plin2 colocalizing with Alexa647-Dextran as shown in D. Truncated violin plots with median and quartiles are shown. (F) Pearson’s correlation coefficient values quantify the levels of GFP colocalized with mCherry-PLIN2 as shown in D. Truncated violin plots with median and quartiles are shown. (G) CLEM images of AML12 cells expressing mCherry-GFP-PLIN2 and loading with Alexa647-Dextran as shown in Figure S2A. Box framed regions are enlarged (Zoom). Yellow arrowheads indicate the contact sites of the lysosomes and LD. SEM, scanning electron microscopy. Scale bars: 10 µm (top panels), 2.5 µm (magnified panels). (H) Z-series SEM images of direct uptake of LD into lysosomes in AML12 cells expressing mCherry-GFP-PLIN2 and loading with Alexa647-Dextran. Yellow arrowheads indicate the contact sites of the lysosomes and LD. (I-K) Reporter processing assay. AML12 cells were overexpressed with mCherry-GFP-PLIN2 and knocked down with *Lamp2b* (I). Levels of GFP (J) and mCherry-GFP-PLIN2 (K) were quantified. Data are presented as mean ± SD (n = 6). *; nonspecific band. P-values are from unpaired t test (F), Dunnett’s multiple comparisons test (A) or Tukey’s multiple comparisons test (B, E, J and K).

To investigate whether microlipophagy occurs at the contact site between LDs and lysosomes, correlative light and electron microscopy (CLEM) was performed. Direct uptake of LD into lysosomes was observed at the contact sites of the lysosomes and LD (Figure 3G, 3H, S2C, S3A, S3B, Video S1), indicating that LAMP2B enhanced lipid hydrolysis through microlipophagy.

Next, we determined whether LAMP2B is involved in the degradation of proteins localized on the LD surface using reporter processing assay with an mCherry-GFP-Plin2 reporter. LAMP2B KD inhibited the cleavage of GFP in cells with or without added OA (Figure 3I and J). Degradation of full-length mCherry-GFP-Plin2 was also inhibited upon KD of LAMP2B (Figure 3I and K). These results suggest that LAMP2B also mediates the degradation of LD surface proteins during LD hydrolysis.

### Pathway involved in LAMP2B-mediated lipid hydrolysis is ESCRT-dependent microlipophagy, not macroautophagy

Next, we investigated whether LAMP2B-mediated lipid hydrolysis is independent of macroautophagy. Macroautophagic flux assay, which analyzes the effect of CQ on LC3-II levels, was performed to determine whether LAMP2B-mediated lipid hydrolysis differs from macroautophagy, and the results showed that macroautophagic flux was not significantly affected by LAMP2B KD in AML12 cells (Figure 4A, 4B). Similarly, LAMP2B KD or overexpression did not significantly affect macroautophagic flux in HeLa cells (Figure S4A-D). Additionally, treatment of cells with 3-methyladenine, an inhibitor of macroautophagy, upregulated cellular lipid levels (TAG and cholesteryl ester) and subsequent LAMP2B overexpression attenuated LD accumulation (Figure S4E). Furthermore, we used *Atg5* knockout (KO) mouse embryonic fibroblasts (MEFs) with inhibited macroautophagy activity^28^ and observed that KD of LAMP2B induced LD accumulation and decreased the levels of lipid hydrolysis (Figure 4D, 4E, S4F). Overexpression of LAMP2B in *Atg5* KO MEFs increased lipid hydrolysis (Figure 4F). These results confirmed that LAMP2B-mediated lysosomal lipid hydrolysis differs from macroautophagy.

**Figure 4.**
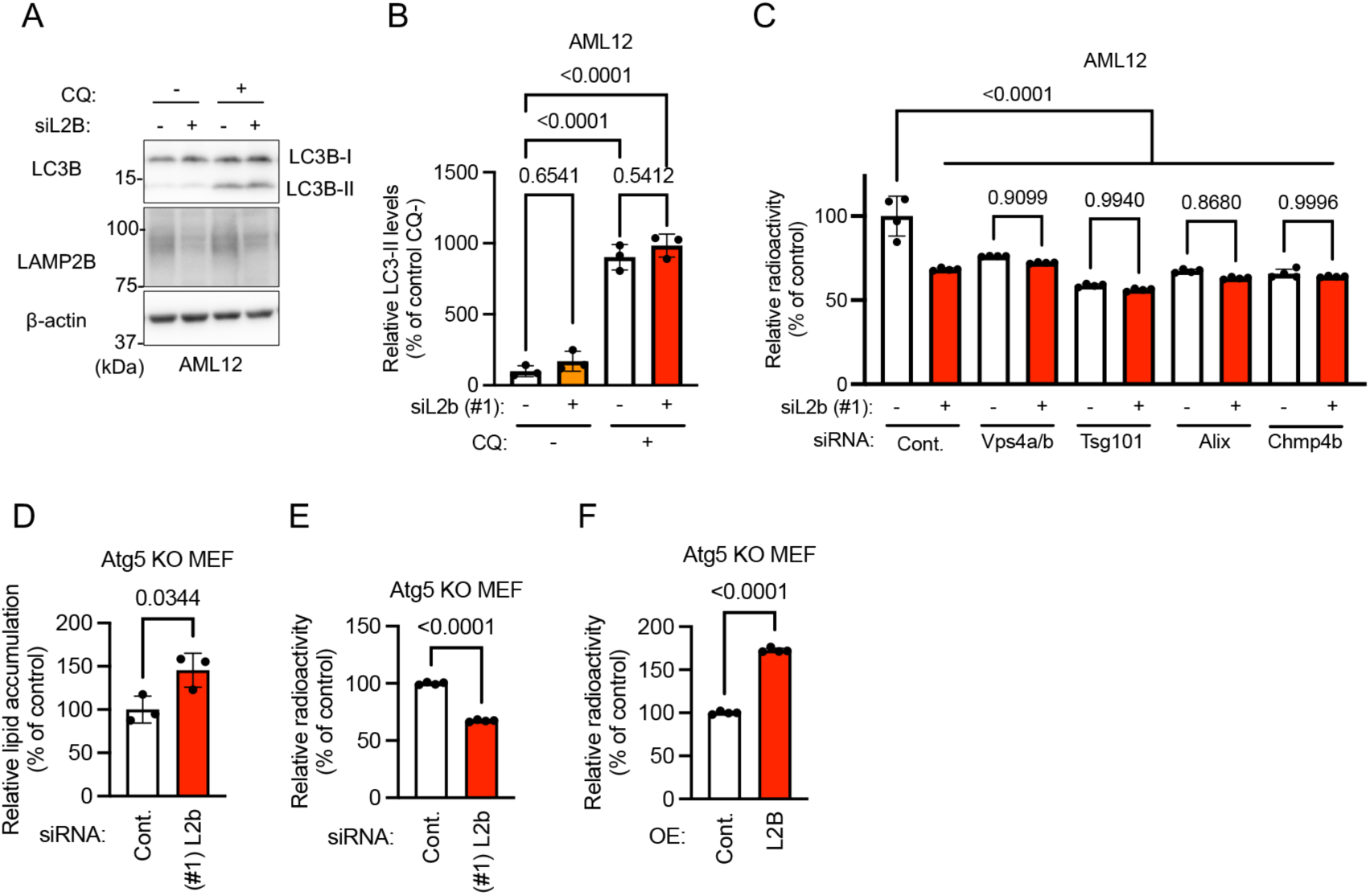
LAMP2B-mediated lipid hydrolysis differs from macroautophagy and is ESCRT-dependent macrolipophagy. (A and B) Macroautophagic flux assay of AML12 cells transfected with *Lamp2b* or Cont. siRNA. The levels of LC3-II were analyzed by immunoblotting (A), and quantified (B). Data are presented as mean ± SD (n = 3). (C) AML12 cells were transfected with *Lamp2b* and *Vps4a/b*, *Tsg101*, *Alix*, *or Chmp4b*-siRNA and lipid hydrolysis level was measured. Data are presented as mean ± SD (n = 4). (D) *Atg5*^−/−^ MEFs transfected with *Lamp2b*- or *EGFP*-#1 siRNA were stained with Oil Rsed O, and dye retention was measured using a spectrophotometer. Data are presented as mean ± SD (n = 3). (E) *Atg5*^−/−^ MEFs were transfected with *Lamp2b*- or *EGFP*-#1 siRNA and lipid hydrolysis level was measured. Data are presented as mean ± SD (n = 4). (F) *Atg5*^−/−^ MEFs were overexpressed with LAMP2B, and lipid hydrolysis level was determined. Data are presented as mean ± SD (n = 4). P-values are from Tukey’s multiple comparisons test (B and C), unpaired t test (D, E, and F).

Recently, it has been reported that the degradation of STING-positive vesicles via microautophagy requires endosomal sorting complexes required for transport (ESCRT) in mammalian cells.^29^ Therefore, we investigated whether LAMP2B-mediated microautophagy requires ESCRT machinery. We examined the effect of double KD of LAMP2B and various ESCRT factors (TSG101 subunit of ESCRT-I, ESCRT-associated protein ALIX (PDCD6IP), CHMP4B subunit of ESCRT-III, or VPS4A and VPS4B for disassembly the ESCRT complexes). KD of VPS4A/B, TSG101, ALIX, or CHMP4B alone decreased lipid hydrolysis indicating the existence of ESCRT-dependent lipid hydrolysis in AML12 cells (Figure 4C, S4G). Intriguingly, combined KD of LAMP2B and ESCRT factors did not enhance lipid hydrolysis compared with KD of LAMP2B or ESCRT factors alone (Figure 4C, S4G). These results suggest that LAMP2B-mediated lipid hydrolysis is conducted via ESCRT-dependent microlipophagy.

### LAMP2B-mediated lipid hydrolysis requires lysosomal localization and interaction of LAMP2B with phospholipid molecules

Next, we assessed the significance of the lysosomal localization of LAMP2B, using the mutant L410A LAMP2B,^30^ which is mutated in the lysosomal targeting signal ‘GYXXФ’ at the C-terminal and localizes to the plasma membrane instead of lysosomes, (Figure 5A and 5B) while interacting with phospholipids (Figure 5C). The results showed that overexpression of L410A LAMP2B did not affect lipid hydrolysis or LD accumulation (Figure 5D, 5E, and S5), indicating that LAMP2B-mediated lipid hydrolysis required the lysosomal localization of LAMP2B.

**Figure 5.**
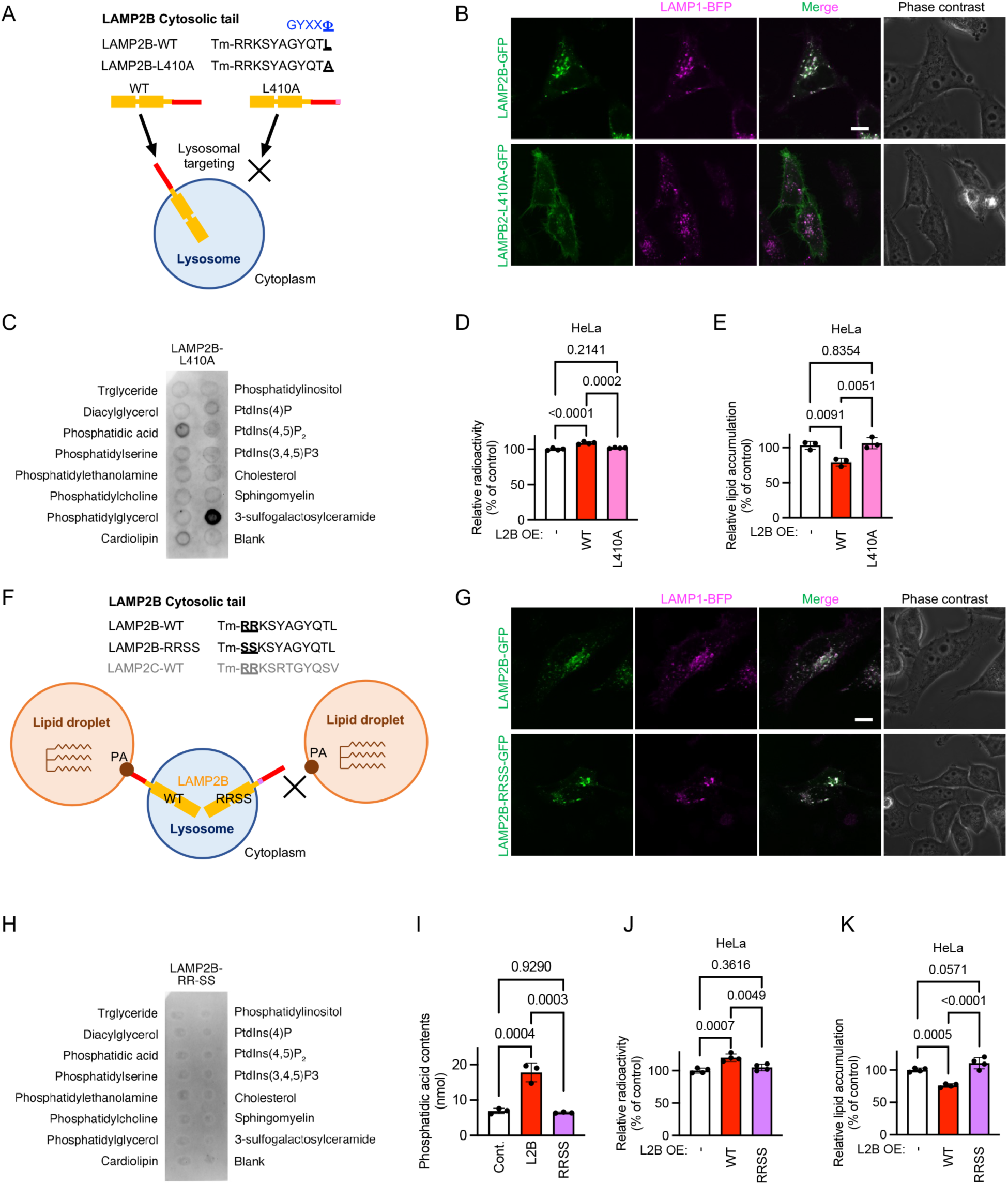
LAMP2B-mediated lipid hydrolysis requires lysosomal localization and interaction of LAMP2B with phospholipid molecules. (A) Schematic of cellular localization of LAMP2B-WT and -leucine to alanine (L410A) mutant. (B) Fluorescence images of HeLa cells expressing LAMP2B WT or L410A–GFP and LAMP1-BFP. Scale bar: 10 µm. (C) Membrane lipid binding assay of LAMP2B-L410A cytosolic tails. (D) Lipid hydrolysis level of HeLa cells transfected with indicated expression vectors. Data are presented as mean ± SD (n = 4). (E) Oil red O-based quantification of LD accumulation in HeLa cells transfected with indicated expression vectors. Data are presented as mean ± SD (n = 4). (F) Schematic of LAMP2B arginine-arginine to serine-serine (RRSS) mutant. (G) Fluorescence images of HeLa cells expressing LAMP2B WT or RRSS–GFP and LAMP1-BFP. Scale bar: 10 µm. (H) Membrane lipid binding assay of LAMP2B-RRSS cytosolic tails. (I) The interaction between LAMP2B-WT or -RRSS cytosolic tails and PA was examined by pull-down assay. Data are presented as mean ± SD (n = 3). (J) Lipid hydrolysis level of HeLa cells transfected with indicated expression vectors. Data are presented as mean ± SD (n = 4). (K) Oil red O-based quantification of LD accumulation in HeLa cells transfected with indicated expression vectors. Data are presented as mean ± SD (n = 4). P-values are from Tukey’s multiple comparisons test (D, E, I, J, and K).

To examine the effect of LAMP2B interaction with PA on LAMP2B-mediated lipid hydrolysis, we constructed a mutant LAMP2B that localizes to lysosomes and does not interact with PA, as well as investigated the effect of the mutation on LAMP2B-mediated lipid hydrolysis. We previously reported that the arginine residues in the cytosolic regions of LAMP2C are required for the interaction between LAMP2C and nucleic acids.^18^ These residues are conserved within LAMP2B (Figure 5F), and RNA and PA competitively bind to LAMP2B (Figure 1B). Therefore, we hypothesized that the arginine residues in the cytosolic regions of LAMP2B are required for the interaction between LAMP2B and PA (Figure 5F). We constructed a mutant in which two arginines were substituted with serines (RRSS mutant). LAMP2B-RRSS mutants colocalized with LAMP1-positive lysosomes, similar to LAMP2B wild-type (WT) cells (Figure 5G). Lipid array analysis using LAMP2B-RRSS mutants revealed that the mutants did not bind to PA (Figure 5H) (control: Figure 1D), which was further confirmed by a pull-down assay (Figure 5I). The RRSS mutant did not promote lipid hydrolysis or LD accumulation (Figure 5J, 5K, and S5). Overall, these findings indicated that interaction between LAMP2B and PA is required for LAMP2B-mediated lipid hydrolysis.

### LAMP2B overexpression prevents high-fat diet-induced obesity *in vivo*

To clarify the physiological and pathophysiological roles of LAMP2B on lipid metabolism *in vivo*, we generated *Lamp2b* -deficient (*Lamp2b*-KO) mice and *Lamp2b* transgenic (*Lamp2b*-Tg) mice that overexpress LAMP2B (Figure S6, S7). First, *Lamp2b*-KO and control WT mice were fed a normal diet (ND) or high-fat diet (HFD). The body weight of *Lamp2b*-KO and WT mice fed ND or HFD was not significantly different (Figure S6B, S6C). The weight of white adipose tissue (WAT) relative to body weight was slightly increased in *Lamp2b*-KO mice compared with that in WT mice (Figure S6D). Given the slight increase in WAT weight in *Lamp2b*-KO, we speculated that the function of LAMP2B was compensated by other lipid hydrolysis pathways, such as cytosolic lipolysis and macrolipophagy. Indeed, there was an increase in LAMP2A expression in the WAT of *Lamp2b*-KO mice (Figure S6E). As earlier stated, LAMP2A induces cytosolic lipolysis and macrolipophagy through CMA-mediated degradation of LD proteins.^10^ Therefore, it is difficult to investigate the physiological role of LAMP2B using the KO method.

Next, we analyzed *Lamp2b*-Tg mice, and confirmed the overexpression of LAMP2B throughout the body (Figure S7B). The lifespan of *Lamp2b*-Tg mice was not significantly different from that of WT mice (Figure S7C). The *Lamp2b*-Tg and control WT mice were fed HFD or ND from 4 weeks of age (Figure 6A and S8A-D). HFD-fed *Lamp2b*-Tg mice had significantly lower body weight than HFD-fed WT mice (Figure 6B and Supplemental Table S1), despite similar feed intake and body temperature (Figure S8E, S8F). There was no significant difference in the body weight of ND-fed *Lamp2b*-Tg mice and WT mice (Figure 6B and Supplemental Table S1). Interestingly, there was no significant difference in the body weight of HFD-fed *Lamp2b*-Tg mice and ND-fed WT mice aged 4–18 weeks (Figure 6B, and Supplemental Table 3).

**Figure 6.**
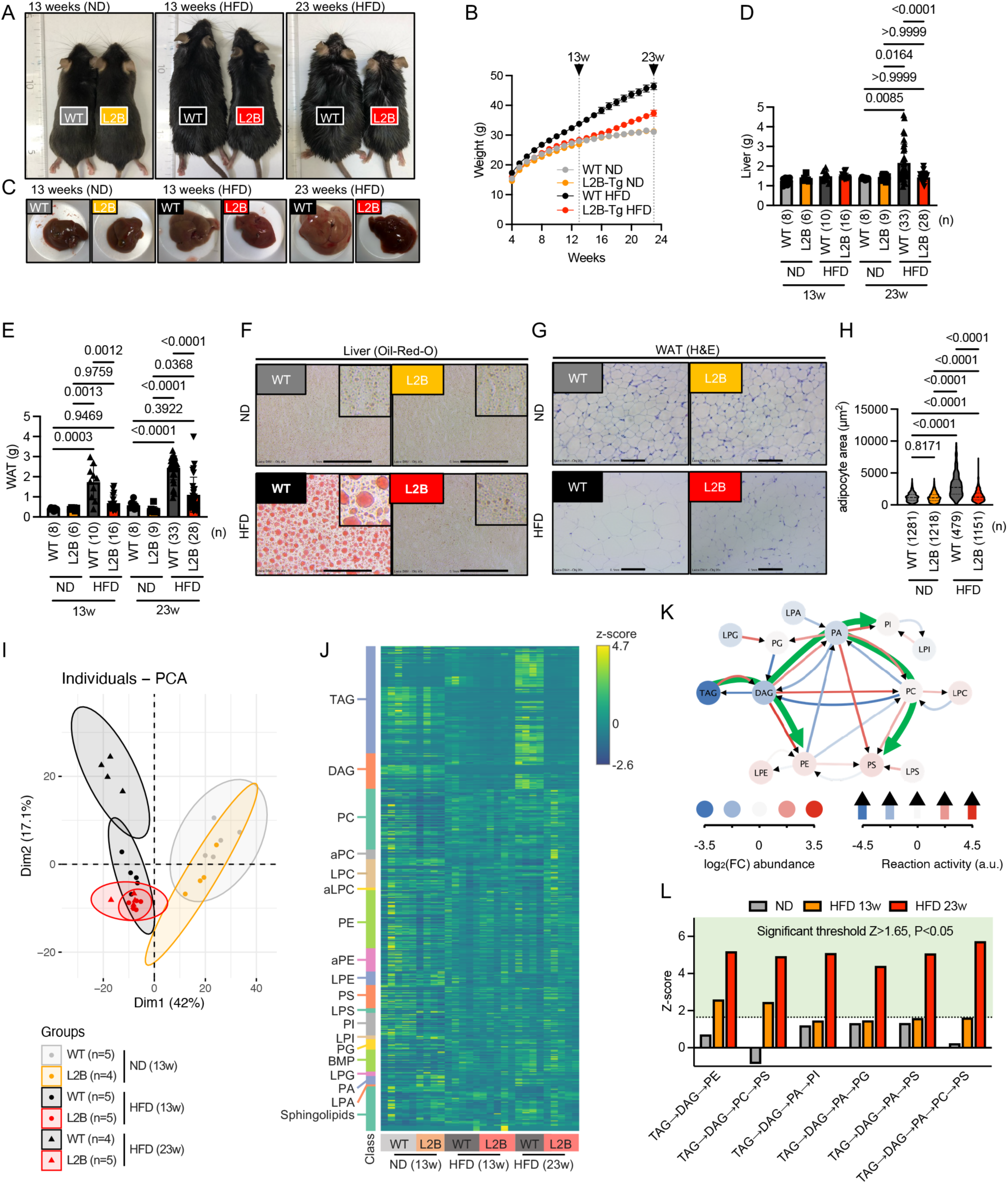
LAMP2B overexpression alters TAG metabolism and prevents HFD-induced obesity. (A) Representative photos of WT and *Lamp2b*-Tg (L2B) mice fed ND for 13 weeks and HFD for 13 or 23 weeks. (B) Body weights of WT and *Lamp2b*-Tg mice fed ND or HFD for the indicated periods. Data are presented as mean ± SEM. The number of samples (n) and p-values were shown in Supplemental Table S1. (C) Representative appearance photos for liver. (D and E) Weights of liver (D) and WAT (E). Data are presented as mean ± SD. (F) Oil red O staining of the liver of WT and *Lamp2b*-Tg mice (scale bar: 0.1 mm). (G) Hematoxylin and eosin staining of WAT of WT and *Lamp2b*-Tg mice (scale bar: 0.1 mm). (H) Quantification of adipocyte area of G. Truncated violin plots with median and quartiles are shown. (I and J) PCA (I) and heatmap visualization (J) of liver lipidome. (K and L) Lipid reaction network analysis in the *Lamp2b*-Tg relative to WT mice fed HFD for 23 weeks. Red and blue circles represent the relative log2-transformed abundances of lipid classes (red: increased, blue: decreased). Red and blue arrows indicate reactions with increased and decreased activity, respectively. Green arrows highlight the major predicted lipid fluxes across the network (K). Increased pathways in the *Lamp2b*-Tg relative to WT mice fed HFD for 23 weeks are illustrated (a dashed line indicates the significance threshold) (L). P-values are from Tukey’s multiple comparisons test (D, E, and H).

Additionally, we analyzed the organ weights in mice aged 13 or 23 weeks. The results showed that HFD-fed mice had increased weights of the liver, WAT, pancreas, brown adipose tissue (BAT), lung, and spleen, but not of the stomach, intestine, quadriceps, heart, kidney and testis (Figure 6C, 6D, 6E, and S8G). We found that the weights of the liver, WAT, pancreas, BAT, and lung in HFD-fed *Lamp2b*-Tg mice decreased compared with those in HFD-fed WT mice, and were not significantly different from those in ND-fed WT mice (Figure 6C, 6D, 6E, and S8G). Furthermore, Oil Red O staining showed increased cellular TAG levels in the liver of WT mice, which was suppressed in HFD-fed *Lamp2b*-Tg mice (Figure 6F). Moreover, H&E staining showed that there was a significant decrease in the size of adipocytes in HFD-fed *Lamp2b*-Tg mice compared with in HFD-fed WT mice (Figure 6G, 6H). These data revealed that LAMP2B prevents HFD-induced obesity *in vivo*.

### Lipidome analysis revealed alteration of TAG metabolism in HFD-fed *Lamp2b*-Tg mice

To determine whether lipid hydrolysis is also promoted in HFD-fed *Lamp2b*-Tg mice, lipidome analyses were performed for all experimental mice (ND- or HFD-fed WT or *Lamp2b*-Tg mice). We identified 22 major classes of lipids, comprising 612 molecular species, in the liver using LC-MS/MS (Table S2). Upon performing principal component analysis (PCA), we observed that principal component (PC) 1 represented approximately 42%, and PC2 represented approximately 17.1% of the total variation (Figure 6I). PC1 separated the profiles by the differences in feed. PC2 separated the profiles by genotype (Figure 6I); in particular, the HFD-fed WT and *Lamp2b*-Tg mice profiles were clearly separated (Figure 6I). Next, we created a heatmap based on the lipidome analysis (Figure 6J). We observed that TAG and diacylglycerol (DAG) levels increased in HFD-fed WT mice but not in HFD-fed *Lamp2b*-Tg mice (Figure 6J). Furthermore, multiple molecular species of TAG and DAG significantly increased in HFD-fed WT mice, but not in HFD-fed *Lamp2b*-Tg mice (Figure S9, S10, and Table S2). Finally, we determined whether lipid metabolism was affected in *Lamp2b*-Tg mice. We used a lipid reaction network analysis method^31^ to assess the activities of glycerolipid and glycerophospholipid pathways, depending on the metabolite measurements conducted via lipidomics (Figure 6K). Six lipid pathways that are involved in the degradation of TAG to phospholipids significantly increased in *Lamp2b*-Tg mice compared with in WT mice (TAG-DAG-phosphatidylethanolamine (PE), TAG-DAG-phosphatidylcholine (PC)-phosphatidylserine (PS), TAG-DAG-PA-phosphatidylinositol (PI), TAG-DAG-PA-phosphatidylglycerol (PG), TAG-DAG-PA-PS, TAG-DAG-PA-PC-PS); meanwhile, five lipid pathways significantly decreased in *Lamp2b*-Tg mice compared with in WT mice (PC-DAG-TAG, PG-DAG-TAG, PC-PA-DAG-TAG, PE-PA-DAG-TAG, lysophosphatidic acid (LPA)-PA-DAG-TAG) (Figure 6L and S11A).

We then measured serum TAG and total cholesterol (T-CHO) using the GPO-HMMPS/glycerol blanking method and the cholesterol oxidase-HMMPS method, respectively. While TAG levels were not affected by HFD (Figure S11B), T-CHO levels were significantly higher in both WT and *Lamp2b*-Tg mice over their ND-fed counterparts (Figure S11C).

Lipidome analysis of the serum showed 24 major classes of lipids composed of 636 molecular species (Table S3). PCA revealed that PC1 represents approximately 40.9% and PC2 represents approximately 13.1% of the total variation, with a clear separation in the profiles of HFD- and ND-fed mice, along with WT and *Lamp2b*-Tg mice fed HFD for 23 weeks (Figure S11D). Next, we created a heatmap based on lipidome analysis and inferred that PE levels increased in HFD-fed WT mice, which further increased in HFD-fed *Lamp2b*-Tg mice (Figure S11E). Further, we observed that total levels of PE significantly increased in HFD-fed *Lamp2b*-Tg mice compared with in ND-fed WT mice (Figure S11F). Given that PE is among the final products of the upregulated pathways in the liver lipid reaction network analysis (Figure 6L), the elevation of PE in the serum may be due to increase in TAG degradation in the liver of *Lamp2b*-Tg mice.

### LAMP2B overexpression prevents HFD-induced diabetes and insulin resistance *in vivo*

We investigated the effect of LAMP2B overexpression on HFD-induced diabetes and insulin resistance. The fasting blood glucose level of HFD-fed WT mice was significantly higher than that of ND-fed WT mice (Figure 7A); meanwhile, the blood glucose level of HFD-fed *Lamp2b*-Tg mice fed was significantly lower than that of HFD-fed WT mice (Figure 7A). Notably, the blood glucose level of HFD-fed *Lamp2b*-Tg mice was not significantly different from that of ND-fed WT mice (Figure 7A).

**Figure 7.**
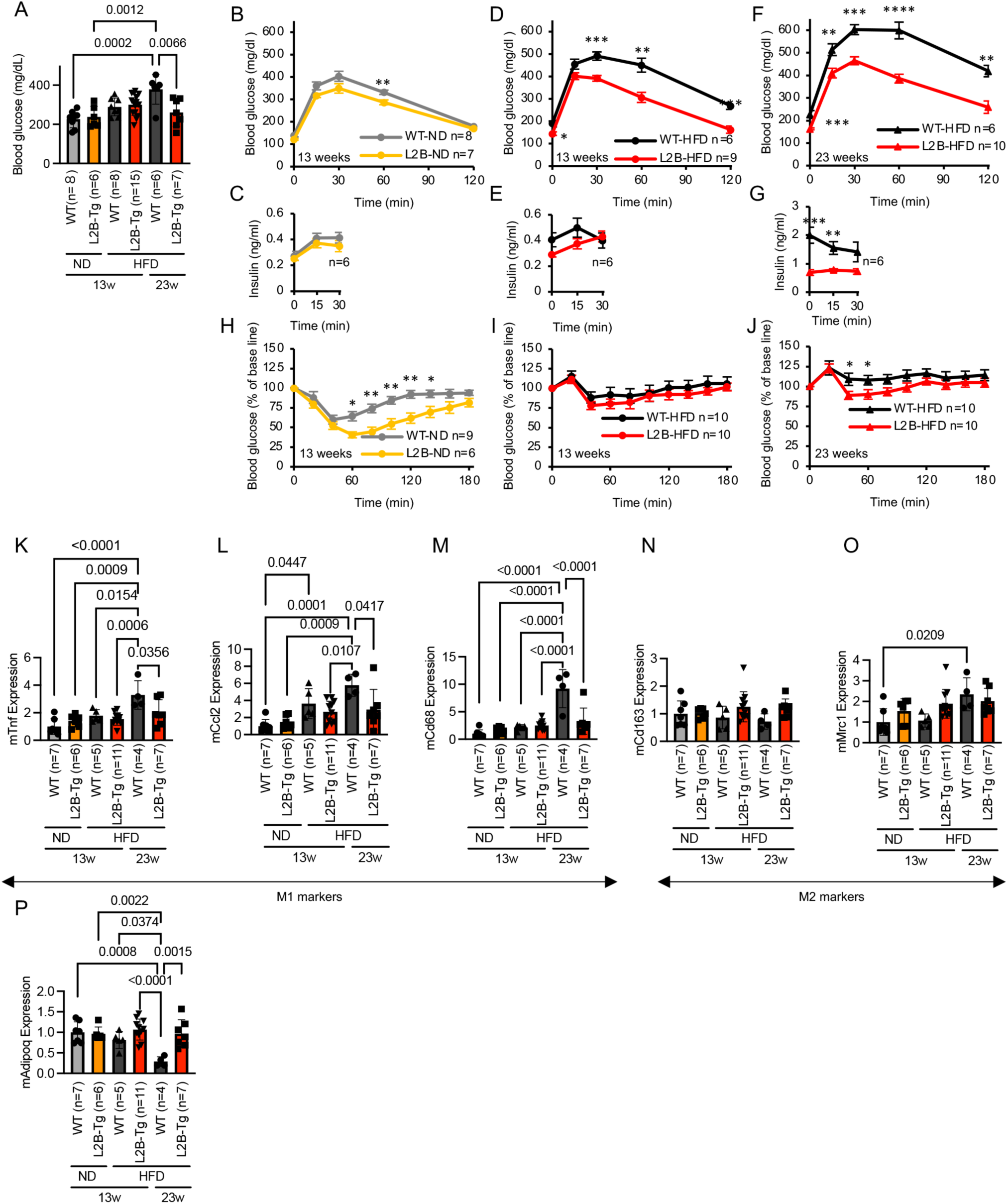
LAMP2B prevents HFD-induced diabetes, insulin resistance, and inflammation in HFD-fed mice. (A) The blood glucose levels in WT and *Lamp2b*-Tg mice. Data are presented as mean ± SD. (B, D, F) Glucose tolerance test using 16 h fasted WT and *Lamp2b*-Tg mice fed ND for 13 weeks (B), HFD for 13 weeks (D) or 23 weeks (F). Data are presented as mean ± SEM. \**p* < 0.05. \*\**p* < 0.01. \*\*\**p* < 0.001. \*\*\*\**p* < 0.0001. (C, E, G) Glucose-stimulated insulin secretion under the conditions in (B, D, F). Data are presented as mean ± SEM. \*\**p* < 0.01. \*\*\**p* < 0.001. (H-J) Insulin tolerance test using 4–6 h fasted WT and *Lamp2b*-Tg mice fed ND for 13 weeks (H), HFD for 13 weeks (I) or 23 weeks (J). Data are presented as mean ± SEM. \**p* < 0.05. \*\**p* < 0.01. (K-P) Expression of M1(*Tnf*: K, *Ccl2*: L, *Cd68*: M) or M2 (*Cd163*: N, *Mrc1*: O) macrophage markers or *Adipoq* (P) in the WAT of WT and *Lamp2b*-Tg mice, with their expression in WT mice fed ND regarded as 1. Data are represented as mean ± SD. P-values are from unpaired t test (B-J), Tukey’s multiple comparisons test (A and K-P).

Next, we performed glucose tolerance test and observed a higher glucose tolerance in ND-fed *Lamp2b*-Tg mice than in WT mice (Figure 7B, 7C). Additionally, HFD-induced glucose intolerance markedly improved in *Lamp2b*-Tg mice (Figures 7D–7G). The serum insulin levels of HFD-fed *Lamp2b*-Tg mice were significantly lower than those of WT mice (Figure 7G), suggesting improved insulin resistance in *Lamp2b*-Tg mice. Furthermore, insulin tolerance test showed that the insulin sensitivity of ND-fed *Lamp2b*-Tg mice was higher than that of WT mice (Figure 7H). Additionally, while the insulin resistance induced by 13 weeks of HFD was similar to that of WT mice (Figures 7I), insulin resistance was found to be suppressed in *Lamp2b*-Tg mice fed HFD for up to 23 weeks (Figures 7J). These results indicated that LAMP2B improved HFD-induced diabetes and insulin resistance *in vivo*.

### LAMP2B reduces adipose tissue inflammation in HFD-fed mice

Adipose tissue is associated with marked infiltration of macrophages during obesity and secretion of adipokines that mediate insulin resistance, a characteristic feature of obesity and type 2 diabetes^32^. To clarify the effect of LAMP2B on adipose tissue inflammation, we examined the expression of M1 (*Tnf*, *Ccl2*, and *Cd68*) and M2 macrophage markers (*Cd163* and *Mrc1*). There was a marked increase in the expression of *Tnf*, *Ccl2,* and *Cd68* in the WAT of HFD-fed WT mice, indicating increased infiltration of pro-inflammatory macrophages, which was not observed in *Lamp2b*-Tg mice (Figure 7K-7M). Among M2 macrophage markers, the expression of *Cd163* decreased, while that of *Mrc1* was increased in HFD-fed WT mice; these effects were not observed in *Lamp2b*-Tg mice (Figure 7N, 7O). Finally, the expression of WAT-specific *Adipoq* decreased in the WAT of HFD-fed WT mice, but not in *Lamp2b*-Tg mice (Figure 7P). Overall, these findings indicate that LAMP2B plays an anti-inflammatory role in adipose tissue inflammation.

## DISCUSSION

To date, the molecular mechanism underlying microlipophagy in mammals has remained elusive. In this study, we show that LAMP2B binds to PA and facilitates microlipophagy. Among the three LAMP2 variants, the cytoplasmic domain of LAMP2A binds to HSC70 and is required for CMA,^15^ whereas that of LAMP2C directly binds to nucleic acids promoting RNautophagy and DNautophagy.^16–19^ Therefore, LAMP2 variants distinguish among various biomolecules, such as proteins, nucleic acids, and phospholipids, through their unique cytoplasmic domains. We previously showed that the cytoplasmic sequence of LAMP2B binds to purified nucleic acids but not in Trizol-treated brain lysate.^17^ Overexpression of LAMP2C promoted intracellular RNA degradation, whereas overexpression of LAMP2B did not.^17^ Conversely, knockdown of LAMP2C also inhibited lipid hydrolysis (Figure 2B), suggesting that LAMP2C functions in both RNA and LD degradation. Taken together, our results suggest that LAMP2B preferentially binds to phospholipids over RNA to promote LD degradation.

Our results revealed that the basic amino acids (RR) in the cytoplasmic region of LAMP2B are essential for binding to PA and promoting lipid hydrolysis (Figure 5H and I). Given that PA is an acidic phospholipid, the binding is probably an electrostatic interaction. Additionally, LAMP2B bound to other phospholipids, such as PtdIns-(3,4,5)-P3 and 3-Sulfogalactosylceramide (Figure 1D). LAMP1 and LAMP2 are reportedly required for fusion of lysosomes with phagosomes.^33^ Therefore, LAMP2B may also be involved in the degradation of organelles other than LDs through phospholipid binding.

PA is a lipid with a cone-shaped structure capable of modulating membrane curvature.^34^ Furthermore, PA functions as both a hydrogen bond donor and acceptor, exhibiting a propensity for clustering.^35,36^ Upon LD-LD fusion, the CIDEA protein that is localized on the LD membrane interacts with PA, disrupting the integrity of the phospholipid barrier at the LD-LD interface, thereby facilitating the transport of TAG molecules previously incorporated in the LD membrane.^20^ Likewise, it is probable that LAMP2B interacts with PA molecules at the LD-lysosome contact sites, inducing LD curvature and facilitating its entry into the lysosome. Nevertheless, while the transport of TAG molecules incorporated into the LD membrane is feasible during LD-LD fusion due to the surrounding phospholipid monolayer, direct transport of TAG (fusion between lysosomes and LDs) is considered difficult because lysosomes possess a phospholipid bilayer. Indeed, CLEM analysis with the LD reporter (mCherry-GFP-Plin2) confirmed the invagination of LDs into lysosomes (Figure 3G, 3H, S2C, S3A, S3B, Video S1). Additionally, the degradation of mCherry-GFP-Plin2 itself is inhibited by LAMP2B knockdown (Figure 3I-K), suggesting that LAMP2B-dependent microlipophagy involves the engulfment of LD, including the LD membrane and its associated proteins, into the lysosome.

Mechanisms involving the lipidation of ATG8 or ESCRT have recently been reported in mammalian microautophagy.^37^ ATG5 is required for the lipidation of ATG8.^28,38^ In the present study, LAMP2B-dependent lipid hydrolysis was observed in *Atg5* KO cells (Figure 4E, 4F), indicating that ATG5 and the lipidation of ATG8 are dispensable for this process. Conversely, knockdown of ESCRT components (TSG101, ALIX, CHMP4B, VPS4A/B) inhibited LAMP2B-dependent lipid hydrolysis (Figure 4C), indicating that LAMP2B-mediated microautophagy functions in an ESCRT-dependent manner. During ESCRT-dependent multivesicular body formation, cargo ubiquitination is crucial for its recognition by ESCRT-0 and ESCRT-I, which subsequently recruit other ESCRT components.^39,40^ Hence, it is possible that ubiquitination of LD plays a significant role in recruiting ESCRT factors in LAMP2B-dependent microlipophagy.

To investigate the pathophysiological significance of LAMP2B-dependent microlipophagy, we employed two approaches by studying both KO and transgenic (overexpressing) LAMP2B mice. While LAMP2B deficiency did not result in increased body weight in mice, it led to a slight increase in WAT (Figure S6D). Two potential reasons may account for the absence of significant weight gain observed in *Lamp2b* KO mice. First, knocking out LAMP2B resulted in upregulation of LAMP2A expression, which possibly promoted cytosolic lipolysis and compensated for LD degradation. Second, since the cytoplasmic region of LAMP2C interacts not only with nucleic acids but also with phospholipids (Figure 1D),^16,17^ it may have compensated for the role of LAMP2B. We verified the therapeutic effect of LAMP2B-mediated lipid degradation on lifestyle diseases using *Lamp2b*-Tg mice. We show that overexpression of LAMP2B promoted TAG degradation in the liver and suppressed obesity (Figure 6). Furthermore, we showed that overexpression of LAMP2B prevented HFD-induced insulin resistance (Figure 7B-J). Additionally, it suppressed the infiltration of inflammatory macrophages into adipose tissue, a phenomenon observed in diabetes (Figure 7K-M). Therefore, overexpression of LAMP2B could be a potentially useful therapeutic strategy to target lifestyle diseases. However, since the HFD-induced elevation in total cholesterol levels did not improve in *Lamp2b*-Tg mice (Figure S11C), we can conclude that LAMP2B overexpression does not ameliorate hyperlipidemia. Interestingly, when fed an ND, *Lamp2b*-Tg mice showed higher glucose and insulin tolerance than WT mice (Figure 7B, 7C, 7H), indicating that LAMP2B may have a glucose tolerance-enhancing effect even when insulin resistance is not observed.

To the best of our knowledge, this study is the first to show the physiological significance of microlipophagy in mammalian organisms. Unlike on HFD, there was no difference in the WAT, liver, or body weight between WT and *Lamp2b*-Tg mice on ND (Figure 6A-H). Therefore, LAMP2B-dependent microlipophagy may be activated in the presence of excess lipids in mice.

Collectively, our study highlights a novel mechanism of microlipophagy in lipid degradation, suggesting that the induction of microlipophagy could serve as a promising therapeutic approach for lifestyle diseases.

## Supporting information

STAR Methods

## Acknowledgments

We thank Dr. Noboru Mizushima, Yoko Ishida, and Keiko Igarashi (The University of Tokyo) for technical assistance with the 3D-CLEM experiments and Yoko Hara (NCNP) for technical assistance with the animal experiments. This work was supported by Grants-in-Aid for Scientific Research from the Japan Society for the Promotion of Science (24K18376 to R.S.; 15K14483, 16H05146, 16H01211, 19H05710, 22H02827, and 23K24089 to T.K.; JP22H04635 to I.K.-H.; 21H04798 and 24H00601 to T.Y.; 21K08565 and 19H05705 to H.-C.L.-O.; and 17K07124 to K.W.), Grants-in-Aid for JSPS Research Fellows (20J00363 to R.S.), research grants from Takeda Science Foundation (to R.S. and T.K.), Intramural Research Grants for Neurological and Psychiatric Disorders (3-9 to T.K.) from the National Center of Neurology and Psychiatry (Japan), and JST-ERATO (JPMJER1702 to I.K.-H. via Noboru Mizushima). This work was supported by the Strategic Research Program for Brain Sciences (SRPBS) from the Japan Agency for Medical Research and Development (AMED) (JP21wm0425008s0201 to T.Y.)

## Author Contributions

R.S., S.A., and T.K. designed and conceptualized the study. R.S., S.A., K.H., and C.K. performed the molecular, biochemical, and cell biological experiments. R.S. and I.K.-H. performed the CLEM. Y.I.-U. and T.I. generated the *Lamp2b*^−/−^ mice. R.S., S.A., K.H., H.F., and H.K. performed the animal experiments. H.-C.L.-O. and T.Y. performed the lipidome experiments. R.S., H.-C.L.-O., T.Y., T.M., and T.K. analyzed the lipidome data. T.H. and K.W. provided critical resources. R.S. and T.K. wrote the manuscript with input from all authors. T.K. supervised the study.

## Declaration of Interests

The authors declare no competing interests.

## Supplementary Tables

**Table S1.** Statistical analysis of body weights in WT and Lamp2b-Tg mice across different ages (weeks).

**Table S2.** Lipidomics data obtained from liver samples.

**Table S3.** Lipidomics data obtained from serum samples.

**Supplementary Video S1.** Z-stack movie of CLEM.

## Supplementary figures

**Figure S1.**
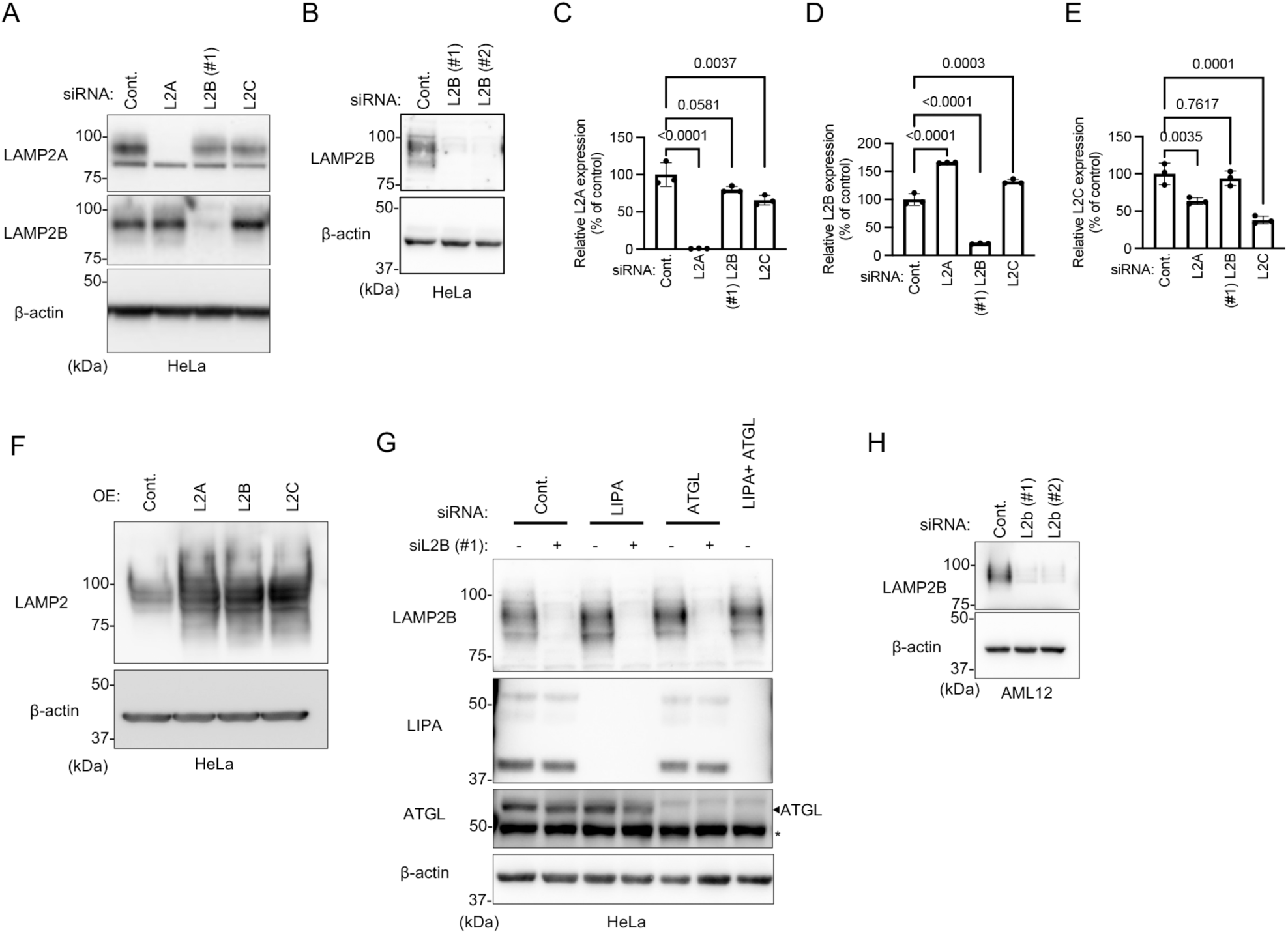
Knockdown and overexpression of LAMP2, related to Figure 2. (A and B) Splice variant-specific knockdown of LAMP2 protein. HeLa cells were transfected with *LAMP2A*, *LAMP2B*, or *LAMP2C*-siRNA, and the expression of the proteins was assessed by immunoblotting. Control cells were transfected with *EGFP*-#2 siRNA. Equal amount of protein was confirmed by immunoblotting for β-actin. (C-E) Splice variant-specific knockdown of LAMP2. HeLa cells were transfected with *LAMP2A*-, *LAMP2B*-, or *LAMP2C*-specific siRNA. Control cells were transfected with *EGFP*-siRNA. Relative expression of each variant was confirmed by real-time quantitative PCR. Data are presented as mean ± SD (n = 3). (F) Overexpression of LAMP2A, LAMP2B, or LAMP2C in HeLa cells. The expression levels of the proteins were assessed by immunoblotting using LAMP2 antibody. Control cells were transfected with empty vector. Equal amount of protein was confirmed by immunoblotting for β-actin. (G) Knockdown of LAMP2B, LIPA and ATGL protein. HeLa cells were transfected with indicated siRNAs, and the expression of the proteins was assessed by immunoblotting. Control cells were transfected with *EGFP*-#2 siRNA. Equal amount of protein was confirmed by immunoblotting for β-actin. *; nonspecific band. (H) Knockdown of LAMP2B protein in AML12 cells transfected with *Lamp2b*-#1 or #2 siRNA was assessed by immunoblotting. Control cells were transfected with *EGFP*-#1 siRNA. Equal amount of protein was confirmed by immunoblotting for β-actin. P-values are from Dunnett’s multiple comparisons test (C-E).

**Figure S2.**
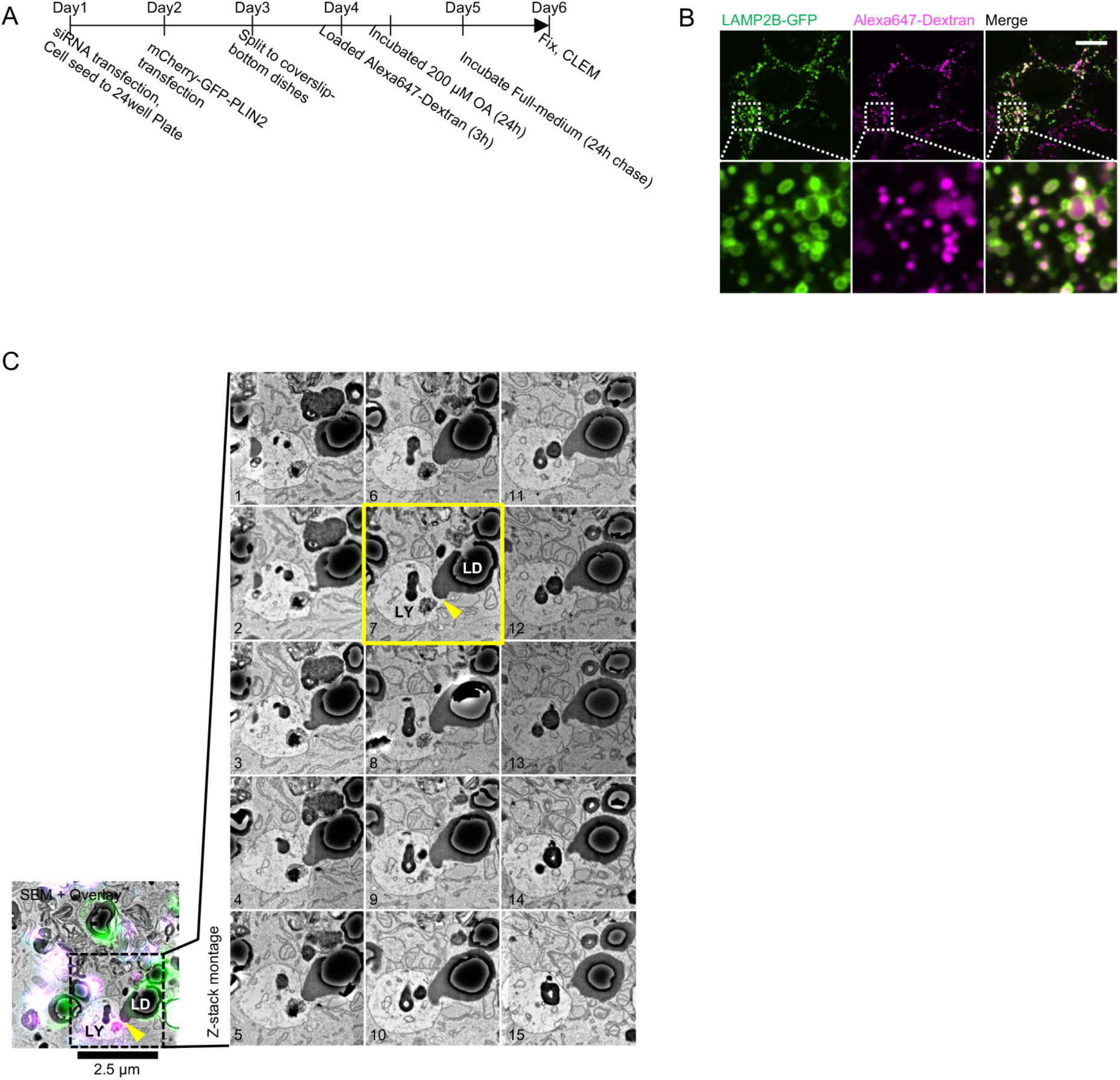
Additional information regarding the study using CLEM, related to Figure 3. (A) Schematic of cell preparation for CLEM. (B) Colocalization of Alexa647-Dextran signals with LAMP2B-GFP-positive lysosomes. AML12 cells overexpressed with LAMP2B-GFP were treated with Alexa647-Dextran for 3 h and incubated for 48 h as shown in Figure. S2A. Scale bar: 10 µm. (C) Full Z-series SEM images of Figure 3H. Yellow arrowheads indicate the contact sites of the lysosomes and LD.

**Figure S3.**
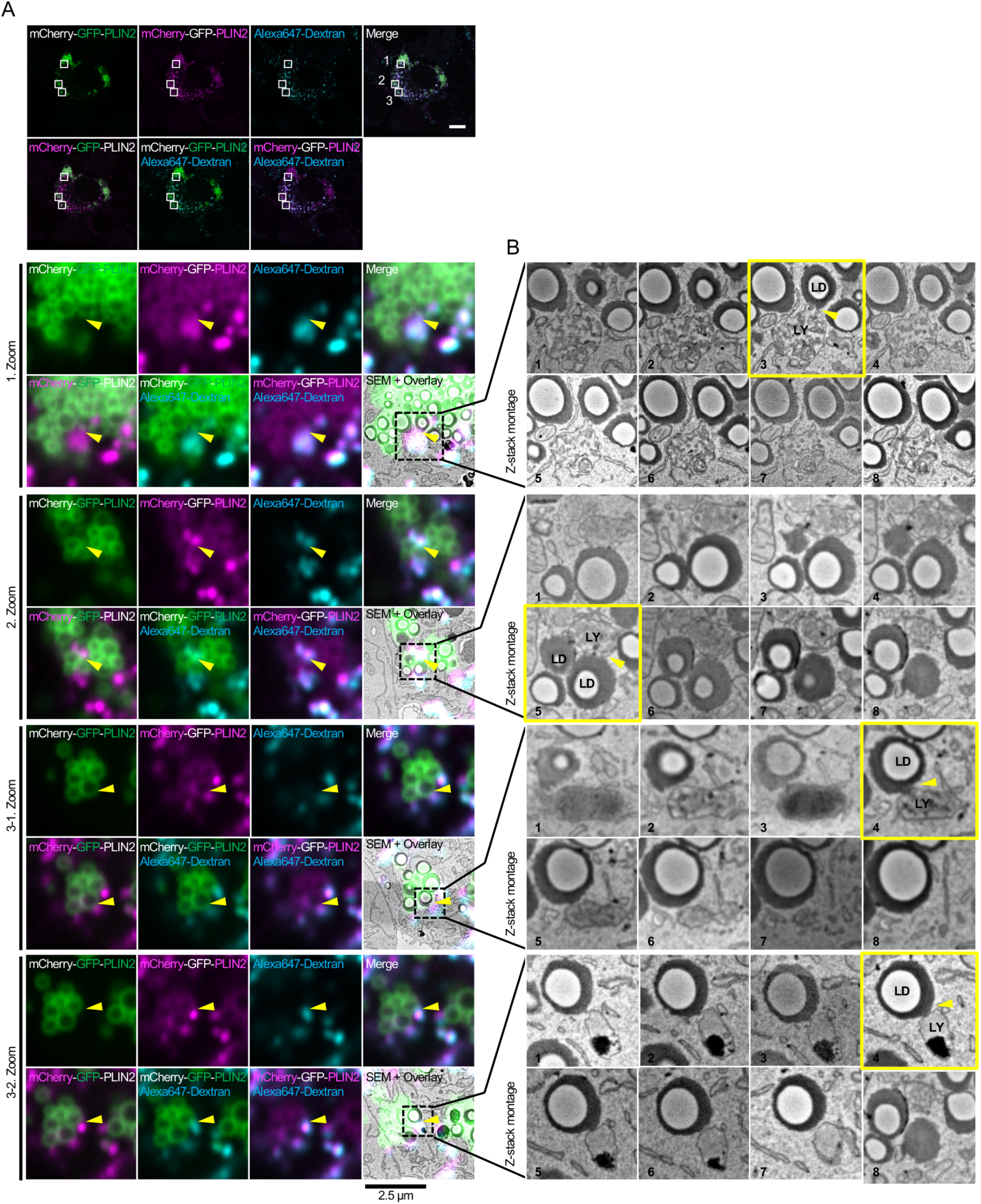
Additional examples of LD-lysosomal interaction using CLEM, related to Figure 3. (A) CLEM images of AML12 cells expressing mCherry-GFP-PLIN2 and loading with Alexa647-Dextran as shown Figure S2A. Box framed regions are enlarged (Zoom). Yellow arrowheads indicate the contact sites of the lysosomes and LD. Scale bars: 10 µm (top panels), 2.5 µm (magnified panels). (B) Z-series SEM images of direct uptake of LD into lysosomes in AML12 cells expressing mCherry-GFP-PLIN2 and loading with Alexa647-Dextran.

**Figure S4.**
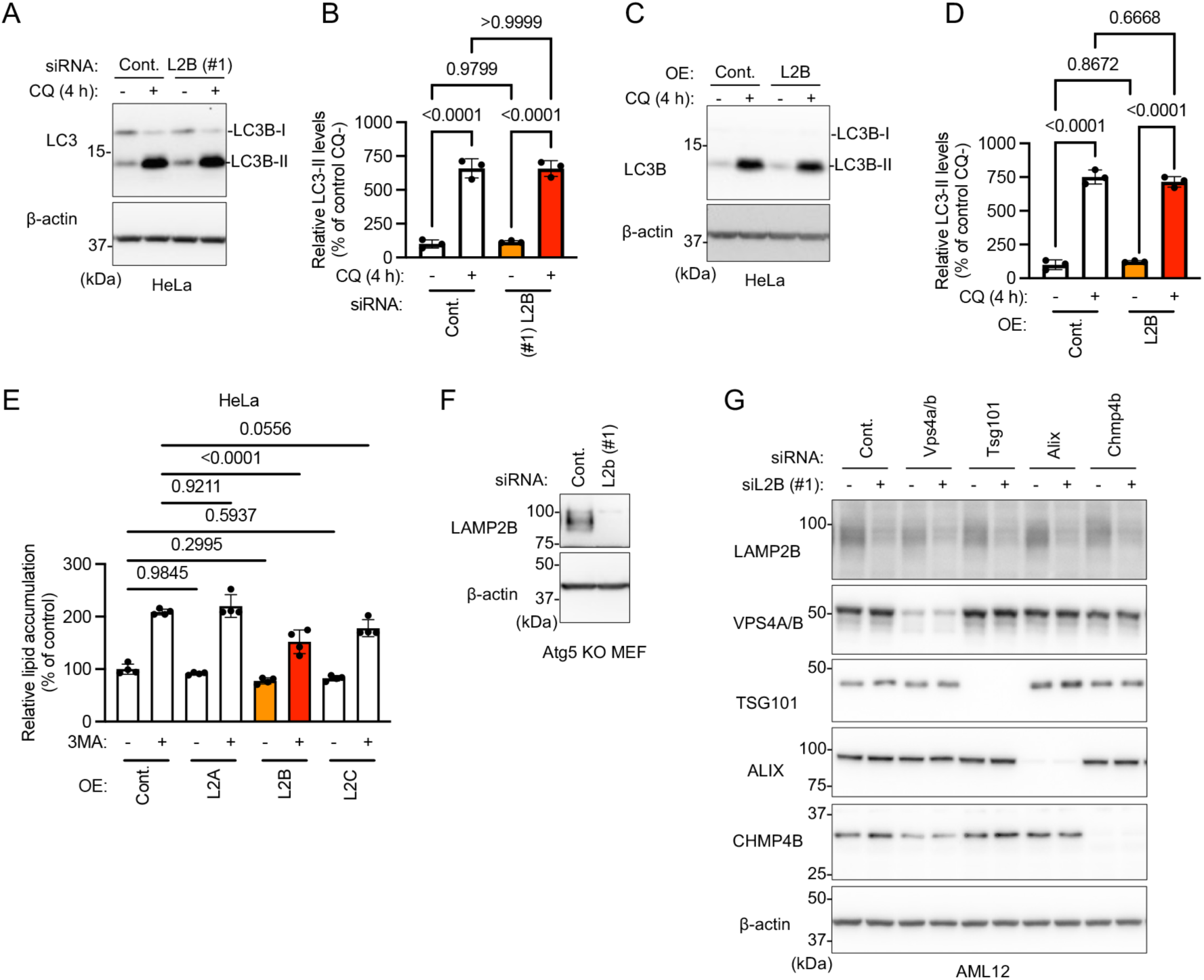
LAMP2B-mediated lipid hydrolysis differs from macroautophagy and is ESCRT-dependent macrolipophagy, related to Figure 4. (A and B) Macroautophagic flux assay of HeLa cells transfected with *LAMP2B* or *EGFP*-#2 siRNA. The levels of LC3-II were analyzed by immunoblotting (A), and quantified (B). Data are presented as mean ± SD (n = 3). (C and D) Macroautophagic flux assay of HeLa cells overexpressed with LAMP2B. LC3-II levels were analyzed by immunoblotting (C) and quantified (D). Data are presented as mean ± SD (n = 3). (E) Effects of LAMP2 overexpression on 3-MA-induced LD accumulation. Data are presented as mean ± SD (n = 4). (F) Knockdown of LAMP2B protein in *Atg5*^−/−^ MEFs transfected with *Lamp2b*-siRNA was assessed by immunoblotting. Control cells were transfected with *EGFP*-siRNA. Equal amount of protein was confirmed by immunoblotting for β-actin. (G) Knockdown of LAMP2B, VPS4A/B, TSG101, ALIX, and CHMP4B protein. AML12 cells were transfected with indicated siRNAs, and the expression of the proteins was assessed by immunoblotting. Control cells were transfected with *EGFP*-#1 siRNA. Equal amount of protein was confirmed by immunoblotting for β-actin. P-values are from Tukey’s multiple comparisons test (B, D, and E).

**Figure S5.**
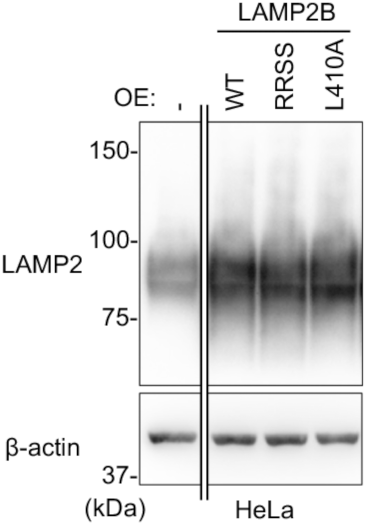
Overexpression of LAMP2B-WT, -L410A or -RRSS mutant protein in HeLa cells, related to Figure 5. The expression levels of the proteins were assessed by immunoblotting using LAMP2 antibody. Control cells were transfected with empty vector. Equal amount of protein was confirmed by immunoblotting for β-actin.

**Figure S6.**
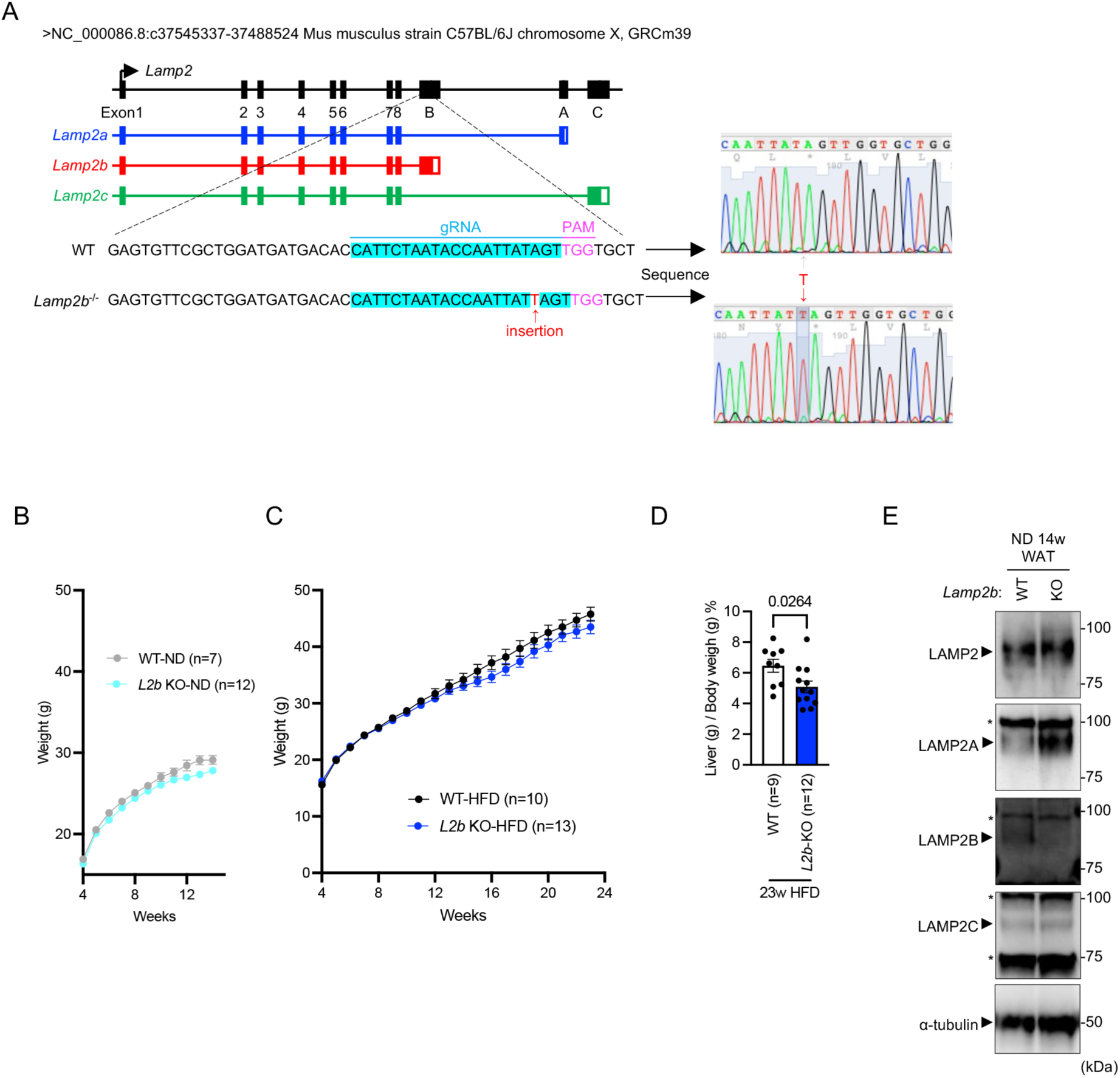
Characterization of *Lamp2b*^−/−^ mice, related to Figure 6. (A) Targeting strategy for *Lamp2b*-specific KO (left panel). The results of the Sanger sequencing of genome of WT and *Lamp2b*^−/−^ mice (right panel). The resulting insertion (T) in the genome of *Lamp2b*^−/−^ mice are in red. (B, C) Body weights of WT and *Lamp2b*^−/−^ mice fed ND (B) or HFD (C) for the indicated periods. Data are represented as mean ± SEM. (D) WAT weights relative to body weights (g/g). Data are presented as mean ± SD. P-values are obtained from unpaired one-tailed t test. (E) The expression levels of LAMP2s proteins in the WAT of the indicated mice were assessed by immunoblotting. *; nonspecific band. Equal amount of protein was confirmed by immunoblotting α-tubulin.

**Figure S7.**
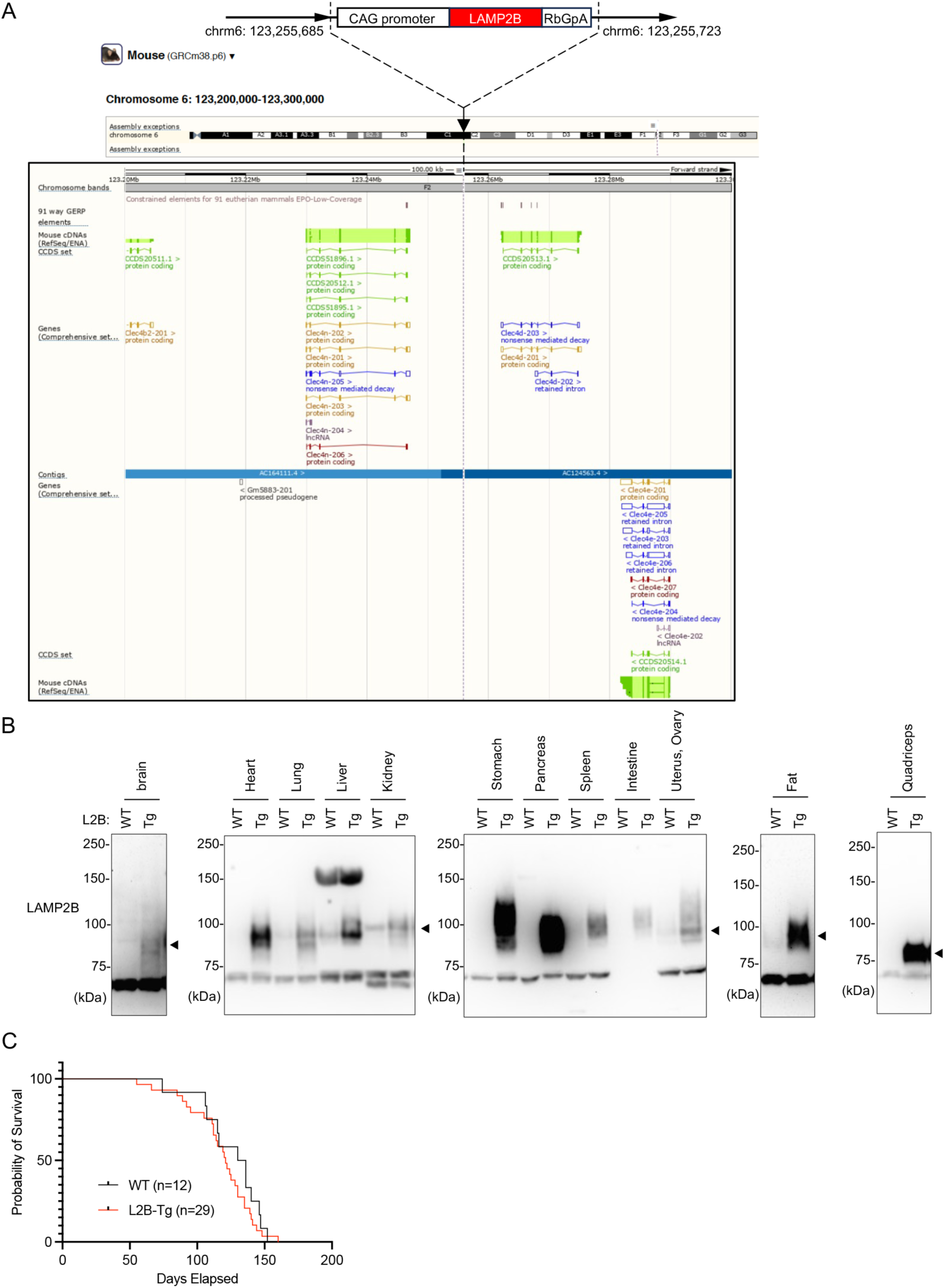
Characterization of *Lamp2b*-Tg mice, related to Figure 6. (A) The insertion point of the *Lamp2b* transgene in *Lamp2b*-Tg mice was identified using whole-genome sequencing. Detailed mapping reveals that the *Lamp2b* transgene has been inserted between 123,255,685 and 123,255,723 base pairs on chromosome 6 (upper panel). Genes proximal to the insertion site of the *Lamp2b* transgene (lower panel). The diagram illustrating the genes was obtained using the Ensembl website (http://www.ensembl.org/). This region is not known to code for any genes, confirming that the insertion of *Lamp2b* gene did not disrupt other genes. (B) Expression levels of LAMP2B protein across various organ tissues in *Lamp2b*-Tg mice (female) were determined by immunoblotting. Specific bands corresponding to LAMP2B are indicated by arrowheads. The protein levels of LAMP2B were elevated in all organs examined in this study. (C) Lifespan of both *Lamp2b*-Tg mice (red) and WT (black) was illustrated with Kaplan-Meier survival curve. There was no significant difference in longevity between the *Lamp2b*-Tg and WT groups, suggesting that overexpression of the LAMP2B does not exhibit toxicity that can affect lifespan.

**Figure S8.**
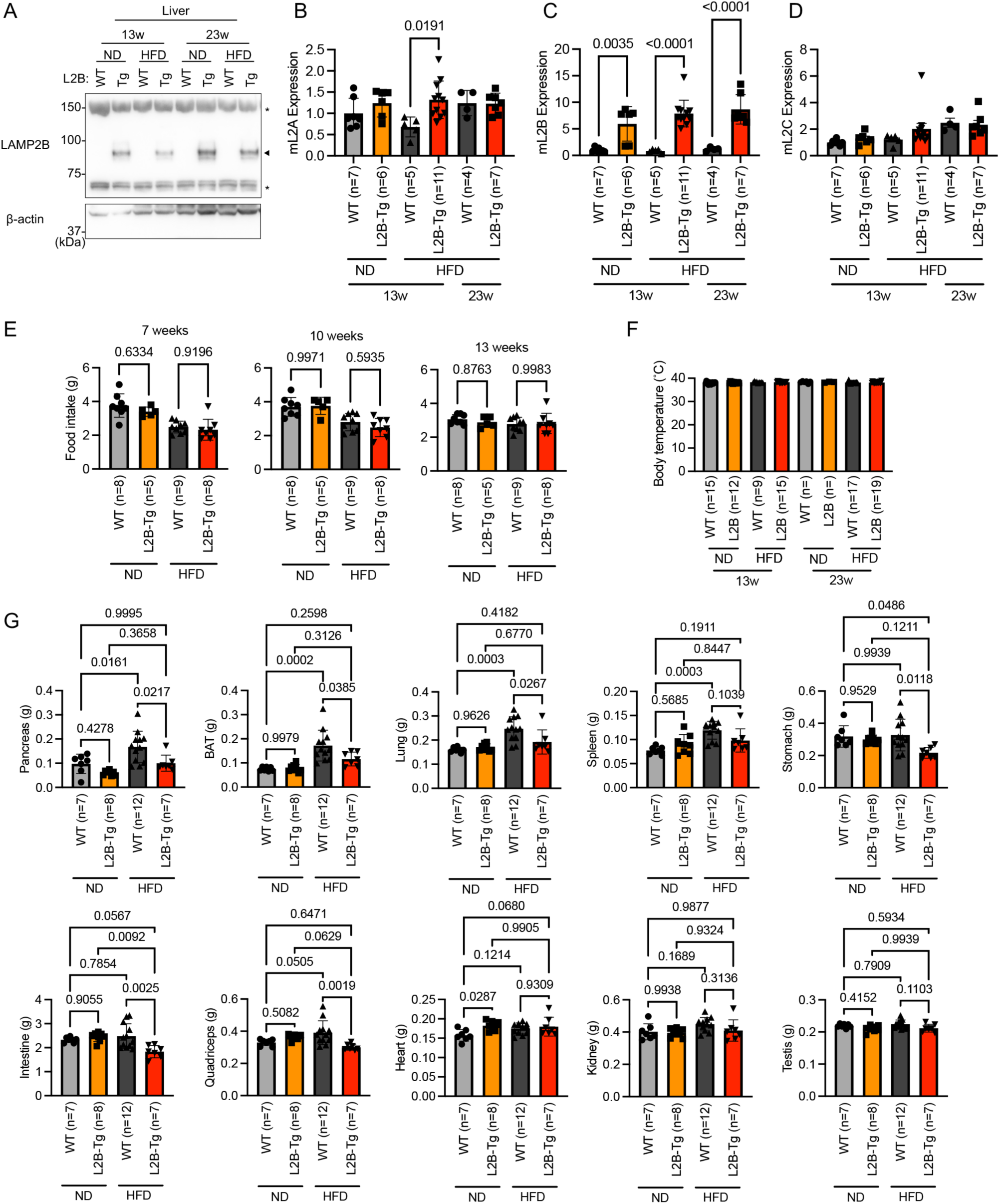
Characterization of *Lamp2b*-Tg mice fed HFD, related to Figure 6. (A) Expression levels of LAMP2B protein in *Lamp2b*-Tg mice fed ND or HFD for 13 or 23 weeks was assessed by immunoblotting. Equal amount of protein was confirmed by immunoblotting for β-actin. *; nonspecific band. (B-D) Relative expression levels of splice variants of *Lamp2* in WAT of *Lamp2b*-Tg mice were assessed by real-time quantitative PCR. Data are presented as mean ± SD. (E) Daily feed intake by WT and *Lamp2b*-Tg mice. Data are presented as mean ± SD. (F) Body temperature of indicated mice. Data are presented as mean ± SD. (G) Weights of indicated organs. Data are presented as mean ± SD. P values are from Tukey’s multiple comparisons test (B-G).

**Figure S9.**
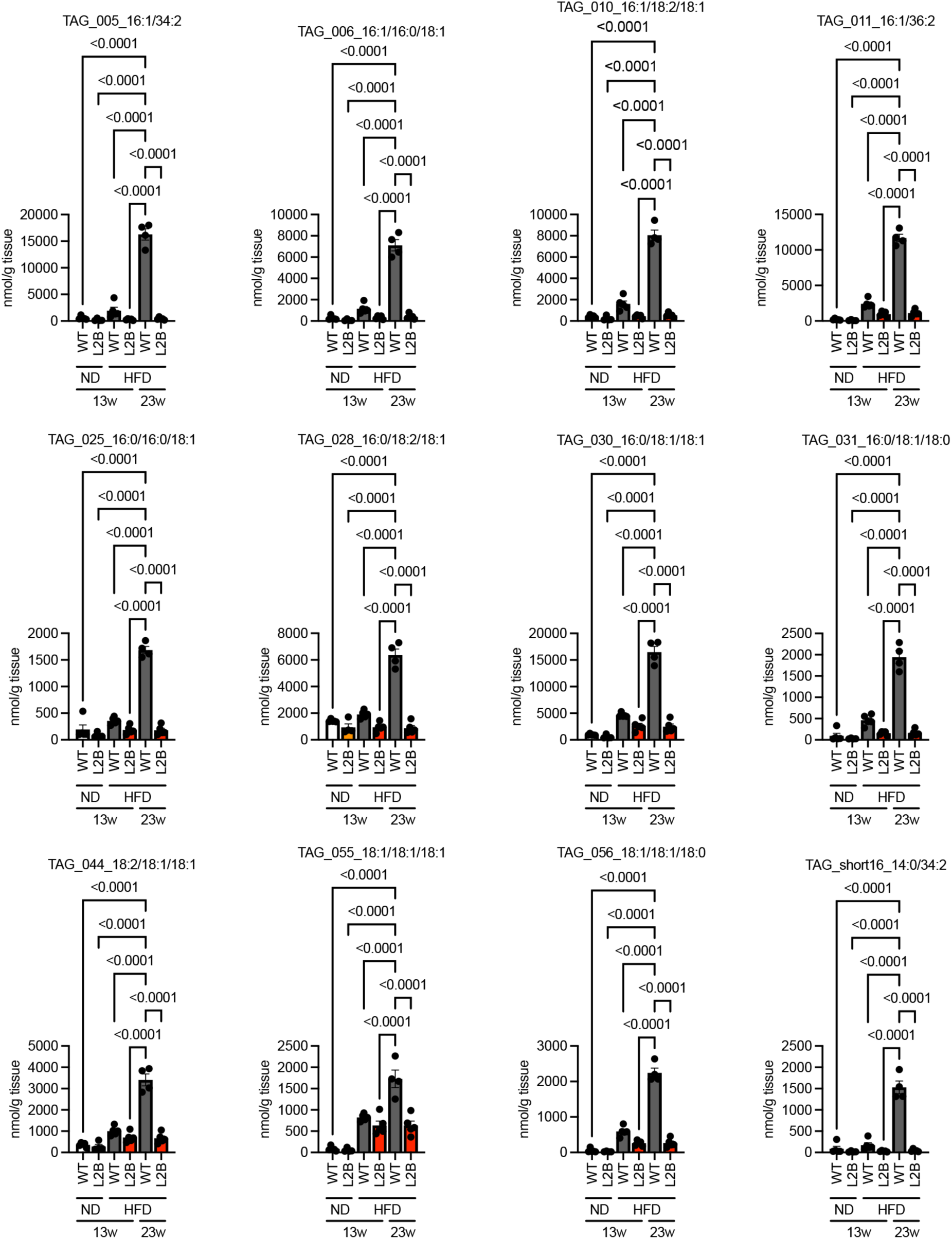
Levels of TAG molecular species in liver lipidome analysis, related to Figure 6. Major TAG molecular species levels, defined as those with mean values above 1500 nmol/g in WT mice fed HFD for 23 weeks. WT mice fed ND for 13 weeks (n = 5), *Lamp2b*-Tg mice fed ND for 13 weeks (n = 4), WT mice fed HFD for 13 weeks (n = 5), *Lamp2b*-Tg mice fed HFD for 13 weeks (n = 5), WT mice fed HFD for 23 weeks (n = 4), and *Lamp2b*-Tg mice fed HFD for 23 weeks (n = 5). Data are presented as mean ± SD. P-values are from Tukey’s multiple comparisons test.

**Figure S10.**
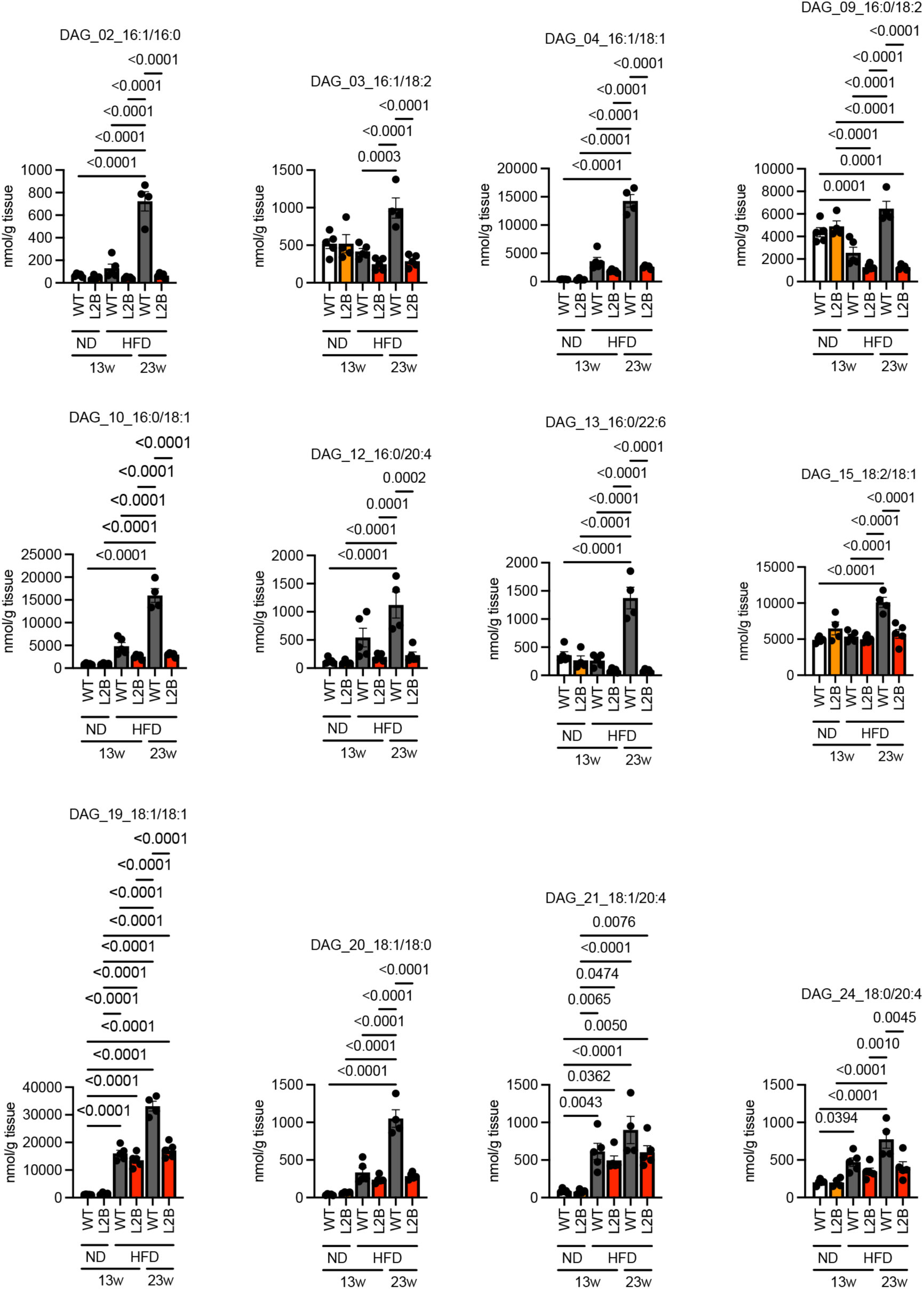
Levels of DAG molecular species in liver lipidome analysis, related to Figure 6. Major DAG molecular species levels, defined as those with mean values above 700 nmol/g in WT mice fed HFD for 23 weeks. WT mice fed ND for 13 weeks (n = 5), *Lamp2b*-Tg mice fed ND for 13 weeks (n = 4), WT mice fed HFD for 13 weeks (n = 5), *Lamp2b*-Tg mice fed HFD for 13 weeks (n = 5), WT mice fed HFD for 23 weeks (n = 4), and *Lamp2b*-Tg mice fed HFD for 23 weeks (n = 5). Data are presented as mean ± SD. P-values are from Tukey’s multiple comparisons test.

**Figure S11.**
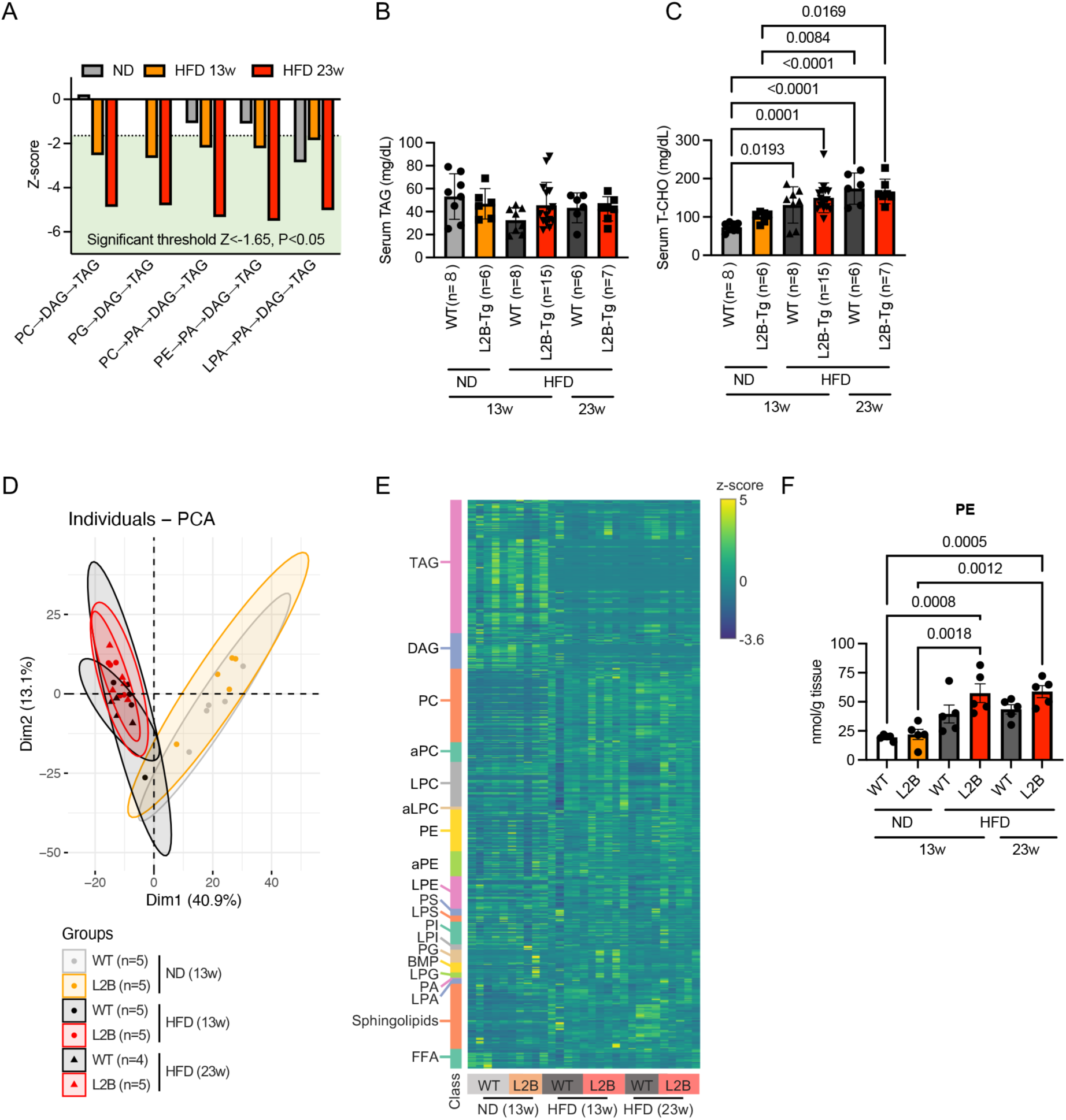
Analysis of lipids in the liver and serum of *Lamp2b*-Tg mice, related to Figure 6. (A) Decreased pathways in the *Lamp2b*-Tg relative to WT mice fed HFD for 23 weeks are illustrated (a dashed line indicates the significance threshold). (B, C) Serum TAG (B) and T-CHO (C) in WT and *Lamp2b*-Tg mice. Data are presented as mean ± SD. (D, E) PCA (D) and heatmap visualization (E) of liver serum. (F) Serum PE in WT and *Lamp2b*-Tg mice. Data are presented as mean ± SD. WT mice fed ND for 13 weeks (n = 5), *Lamp2b*-Tg mice fed ND for 13 weeks (n = 5), WT mice fed HFD for 13 weeks (n = 5), *Lamp2b*-Tg mice fed HFD for 13 weeks (n = 5), WT mice fed HFD for 23 weeks (n = 4), and *Lamp2b*-Tg mice fed HFD for 23 weeks (n = 5). P-values are from Tukey’s multiple comparisons test (B, C, F).

## STAR Methods

### RESOURCE AVAILABILITY

#### Lead contact

Further information and requests for resources and reagents should be directed to the lead contact, Tomohiro Kabuta.

#### Materials availability

All the materials generated in this study are available upon reasonable request to the lead contact.

#### Data and code availability

- This study does not report original code.
- Any additional information required to reanalyze the data reported in the paper is available from the lead contact upon request.

### EXPERIMENTAL MODEL AND STUDY PARTICIPANT DETAILS

#### Cell models

HeLa cells were grown in Dulbecco’s modified Eagle’s medium (DMEM) containing 10% fetal bovine serum (FBS) at 37 °C. The mouse hepatocyte AML12 cell line was obtained from the ATCC (CRL-2254). AML12 cells were grown in Dulbecco’s Modified Eagle Medium/Nutrient Mixture F-12 (DMEM/F-12) supplemented with 10% FBS, insulin-transferrin-sodium selenite media supplement (Sigma), and dexamethasone (40 ng/mL) at 37 °C. *Atg5*-deficient (*Atg5*^−/−^) MEFs were kindly gifted by Dr. Noboru Mizushima (University of Tokyo, Tokyo, Japan) and maintained as previously described.^41^

#### Animal treatment

Male mice were used in the study unless mentioned otherwise. Mice of all genotypes were kept on an ND (CLEA Rodent Diet CE-2; Clea, Japan) under a 12-hour dark/light cycle, except for the experimental subgroup, which were maintained on an HFD (High Fat Diet32; Clea, Japan) for 13 or 23 weeks. All procedures in this study were performed in accordance with the guidelines and were approved by the Animal Investigation Committee of the National Institute of Neuroscience, National Center of Neurology and Psychiatry.

#### *Lamp2b*-deficient mice

*Lamp2b*^−/−^ mice were generated using CRISPR/Cas9 system. The sgRNA sequence against a site in exon 9b (5-ATTCTAATACCAATTATAGT-3) of *Lamp2* was designed using CRISPOR (http://crispor.tefor.net/) and cloned into pX330 plasmid (Addgene, 42230). The plasmid was then directly injected into the pronuclei of B6C3F1 fertilized eggs. The genotypes of the resultant KO mice were confirmed by sequencing analysis. *Lamp2b*-deficient mice were backcrossed with C57BL/6J mice for 7 generations and WT (*Lamp2b*^+/+^) siblings were used in this study. For the immunoblotting analysis, we used 3-4 generation mice.

#### LAMP2B transgenic mice

*Lamp2b* transgenic mice were generated by Unitech. Briefly, a plasmid expressing mouse *Lamp2b* under the control of the CAG (ubiquitous) promoter was constructed. Next, a solution of the linearized plasmid was microinjected into the pronuclei of newly fertilized C57BL/6J mouse eggs. Analysis of the integration site of the transgene using whole-genome sequencing analysis was outsourced to Macrogen.

### METHOD DETAILS

#### Transfection

The cells were transiently transfected with expression vectors using Lipofectamine LTX with Plus Reagent (Life Technologies), according to the manufacturer’s instructions.

#### RNA pull-down assay

Total RNA was purified from mouse brains using TRIzol reagent (Life Technologies), according to the manufacturer’s instructions. RNA pull-down assays were performed as previously described ^19^. Streptavidin Sepharose High-Performance beads (GE Healthcare) were blocked overnight with 3% bovine serum albumin in Dulbecco’s phosphate-buffered saline (PBS) at 4 °C. The blocked beads were incubated at 4 °C for 2 h with 10 µg of purified total RNA and 8 nmol of biotin-conjugated LAMP2B peptides (cytosolic sequence) in the absence or presence of 250 µg of phosphatidic acid, cholesterol, or phosphatidylcholine in 1 mL PBS containing 0.05% Triton-X 100. After incubation, the beads were washed thrice with 0.05% Triton-X 100 in PBS or PBS. RNA was extracted using the TRIzol reagent, and analyzed using agarose gel electrophoresis.

#### Lipid pull-down assay

HeLa cells were lysed in PBS containing 1% Triton-X 100 and centrifuged (15,000 × g, 10 min) at 4 °C, and the supernatant was used as HeLa cell lysate. Streptavidin Sepharose High-Performance beads (GE Healthcare) were blocked as described above. The blocked beads were incubated at 4 °C for 2 h with the HeLa lysate and 8 nmol of biotin-conjugated peptides. After incubation, the beads were washed three times with 0.05% Triton-X 100 in PBS or PBS. Pulled-down PA was extracted and quantified using total phosphatidic acid assay kit (Cayman), according to the manufacturer’s instructions.

#### Membrane lipid binding assay

Biotin-conjugated peptides were synthesized as previously described^17^ Lipid binding assays were performed using a membrane lipid strip obtained from Echelon Bioscience, Inc. Membranes were blocked with 1% fatty acid-free bovine serum albumin (BSA) in PBS for 1 h and probed with the indicated peptides (150 µg/strip) overnight at 4 °C. After overnight incubation, the membranes were washed three times with PBS containing 0.1% Tween-20 (PBS-T), followed by incubation with horseradish peroxidase (HRP)-conjugated streptavidin (Abcam; 1:2,000) in blocking buffer for 1 h. After three washes, the membranes were incubated with SuperSignal West Pico, and chemiluminescence was detected using ImageQuant LAS 4000 (Fujifilm) or FluorChem Chemiluminescence imaging system (Alpha Innotech).

#### RNAi knockdown

Synthetic siRNA oligonucleotides were purchased from Sigma-Aldrich and transfected into cultured cells using Lipofectamine RNAiMAX (Thermo Fisher) for 72 h, according to the manufacturer’s instructions. An siRNA targeting *EGFP*-#1 was used as a non-targeting control in mouse cells. An siRNA targeting *EGFP*-#2 was used as a non-targeting control in human cells. siRNA targeting the universal negative control was used as a non-targeting control when the GFP-tagged plasmid was co-transfected.

#### Lipid hydrolysis measurement

HeLa cells or MEFs were seeded into 24-well plates at a density of 5 × 10^4^ cells per well. In the KD experiments, cells were transfected with the indicated siRNAs at the same time as seeding (reverse-transfection). In the overexpression experiment, the cells were transfected with the indicated expression vectors 24 h after seeding. Exactly 24 h after transfection, the medium was replaced with growth medium supplemented with 400 µM oleic acid, 1 µCi/well [^3^H] oleic acid complex (PerkinElmer), and 0.4% fatty acid-free BSA (Wako) overnight to induce LD formation. The cells were then incubated in a growth medium supplemented with 1% fatty acid-free BSA as a fatty acid acceptor. Triacsin C (Abcam) (5 µM), a known inhibitor of fatty acid acyl coenzyme A ligase, was added to block the re-esterification of the released fatty acids. After 3 h, the radioactivity released into the culture medium was quantified using a liquid scintillation counter (Tri-Carb 3100TR; Packard).

#### Oil red O staining of cultured cells

Cells were grown in 24-well culture plates and transfected with the indicated expression vectors or siRNAs. In some experiments, 200 µM oleic acid (Nacalai Tesque) conjugated with fatty acid-free BSA (WAKO) was added to the culture medium to induce LD formation. At the end of the incubation period, the cells were washed with PBS and fixed using 4% paraformaldehyde in PBS (Wako) for 1 h. After washing with PBS, the cells were rinsed with 60% isopropanol and stained with Oil Red O (Sigma-Aldrich) in 60% isopropanol for 20 min at 25 °C. LDs stained with Oil Red O were observed under a light microscope (BX51; Olympus). To quantify LD accumulation, Oil Red O dye was extracted from cells by adding 100% isopropanol, and the optical density was measured using a spectrophotometer at 490 nm.

#### Confocal imaging

HeLa and AML12 cells cultured on glass-bottom dishes or chamber slides were treated as indicated. Images were captured using a confocal laser microscope (FV1000 or FV10; Olympus). Colocalization of mCherry-GFP-PLIN2 and Alexa647-Dextran was analyzed by Pearson’s correlation coefficient or Manders’ correlation coefficient using ImageJ software.

#### Measurement of lysosomal pH

HeLa cells seeded on 35 mm glass-bottomed dishes were treated with or without 25 mM NH_4_Cl, followed by 15 min of treatment with 1 µM LysoSensor Green DND-189 (Thermo Fisher) 1 h later. The medium was then washed, cells were incubated for 1 h, and lysosomal pH was analyzed as relative fluorescence units using an FV10 confocal microscope (Olympus) and ImageJ software.

#### CLEM

AML12 cells were seeded into 24-well plates at a density of 1 × 10^5^ cells per well and transfected with the indicated siRNAs. The following day, the cells were transfected with mCherry-GFP-Plin2 expression vectors. Exactly 24 h after transfection, the cells were split into no. 1S gridded coverslip-bottom dishes (TCI-3922-035R; Iwaki), precoated with carbon using a vacuum coater, and then coated with gelatin. The following day, the cells were incubated in pre-warmed medium containing 0.1 mg/ mL Alexa Fluor-647-Dextran (Thermofisher) for 3 h at 37 °C. After 3 h, the medium was replaced with growth medium supplemented with 200 µM oleic acid in 0.4% fatty acid-free BSA (Wako) overnight to induce LD formation. Cells were then incubated in growth medium supplemented with 1% fatty acid-free BSA and 5 µM Triacsin C (Abcam) for 24 h. Cells were fixed with 2% paraformaldehyde (26126-54, Nacalai Tesque) and 0.5% glutaraldehyde (G018/1, TAAB) in 0.1 M phosphate buffer (pH 7.4) (LSI Medience Corporation) at room temperature for 1 h. After washing and substitution with phosphate buffer, fluorescent images were obtained using an inverted confocal laser scanning microscope (FV1000) equipped with a PlanApo 60×, 1.4 NA oil immersion objective. After fluorescent image acquisition, the cells were embedded in epoxy resin for electron microscopic observation, and CLEM analysis was performed as previously described.^42^ Briefly, the imaged cells on the gridded-glass were fixed overnight with 2.5% glutaraldehyde in 0.1 M sodium cacodylate buffer at 4 °C, osmicated with 1% osmium tetroxide (3020-4; Nisshin EM) and 1.5% potassium ferrocyanide (161-03742; Wako) in 0.065 M cacodylate buffer for 2 h at 4 °C, en bloc stained with 3% aqueous uranyl acetate at room temperature for 1 h, dehydrated in an ascending series of ethanol, and then embedded in epoxy resin (Epon812, TAAB). The resin block was sectioned using an ultramicrotome (EM UC7, Leica) equipped with a diamond knife (Ultra JUMBO, 35 degree, DiATOME) to cut 25-nm serial sections. The sections were imaged using a scanning electron microscope (JSM7900, JEOL) equipped with Array Tomography Supporter version 1.2.5.0 (System in Frontier) that enables automated imaging. Images were stacked in order using Stacker NEO software version 3.3.4.0 (System in Frontier). The correlation between light and electron microscopic images was performed using ImageJ/FIJI software.

#### Quantitative PCR

Total RNA was isolated using TRIzol reagent (Life Technologies) or TRI Reagent (MRC), and cDNA was synthesized using a PrimeScript RT Reagent Kit with gDNA Eraser (Perfect Real Time) (TaKaRa). PCR amplification was performed using SYBR Premix Ex Taq II (Takara Bio Inc.) or Luna® Universal qPCR Master Mix (BioLabs) on a CFX96 Touch Real-Time PCR Detection System (Bio-Rad). The relative amounts of the target genes were normalized to those of β-actin or GAPDH.

#### Immunoblotting analysis

Protein lysates from cultured cells were separated using SDS-PAGE and transferred onto PVDF membranes (Bio-Rad, 1620177), which were then blocked with 3% BSA (Iwai Chemicals Company) prepared in DPBS or 5% skim milk (Wako) prepared in DPBS containing 0.1% Tween 20 for 30 min at RT. This was followed by overnight incubation with primary antibodies prepared in either of the blocking buffers at 4 °C. The membranes were then washed with 0.1% Tween 20 in DPBS and probed with horseradish peroxidase-conjugated secondary antibodies (Thermo Fisher Scientific). Signals were then visualized using ImmunoStar Zeta (Wako) or ImmunoStar LD (Wako) and detected using FluorChem chemiluminescence imaging system (Alpha Innotech) or FUSION chemiluminescence imaging system (Vilber-Lourmat).

#### Macroautophagic flux assay

The macroautophagic flux assay was performed in HeLa or AML12 cells transfected with indicated siRNAs or plasmids (Figure 4A, S4A, S4C). 48 h after transfection, cells were incubated with or without 50 mM chloroquine (CQ) (Sigma) for 4 h. Then, cell lysates were prepared and subjected to immunoblotting, and the relative levels of LC3B-II and β-actin were quantified. LC3B-II levels were normalized to those of β-actin levels. Macroautophagic flux was calculated by subtracting the normalized LC3B-II levels in the CQ-samples from the levels in the CQ+ samples.

#### Reporter processing assay

In the reporter processing assay employing GFP fusion proteins as substrates, the levels of free GFP indicate autophagic activity.^43,44^ Autophagic substrates expressed as GFP fusion proteins are taken up into lysosomes and degraded, thereby releasing the intact form of GFP, which is detected by immunoblotting. AML12 cells were overexpressed with mCherry-GFP-PLPN2 as described in “Correlative light and electron microscopy (CLEM),” and the levels of free GFP were quantified using immunoblotting.

#### Histological analysis and adipocyte area determination

The WATs were fixed in 10% neutral-buffered formalin, embedded in paraffin, cut into 5 µm sections using a microtome (REM-700; YAMATO), and then mounted on slides. The sections were stained with H&E and photographed using a microscope (Leica DM IL LED) equipped with a digital camera (Leica MC170 HD). The adipocyte area of WAT was analyzed using ImageJ software.

Liver tissues were embedded in Tissue-Tek OCT compound (Sakura, Japan), frozen in liquid nitrogen, cut into 10 µm sections using a cryostat (CM3050 S, Leica Microsystems), and then mounted on slides. The cryosections were stained with Oil Red O and photographed using a microscope (Leica DM IL LED) equipped with a digital camera (Leica MC170 HD).

#### Lipidome analysis of liver and serum samples

Lipids were extracted from frozen tissues (15–60 mg) or sera (5 µl) using the Bligh and Dyer method with internal standards. The organic (lower) phase was transferred to a clean vial and dried under a nitrogen stream. The lipids were resolubilized in a methanol/isopropanol/chloroform mixture (45/45/10, v/v/v) and stored at −80 °C. A portion of the extracted lipids was injected into an ultrahigh-performance liquid chromatography (LC)-electrospray ionization (ESI)–tandem mass spectrometry system (LC-MS/MS). The quantification of FFA was carried out as described previously.^45^ For the quantification of LPA, LPG, LPI, LPS, PA, and PS, LC separation was performed on a 50 × 4.6 mm 5 µm Gemini C18 column (Phenomenex) coupled to a guard column (Gemini; C18; 4 × 3.0 mm; Phenomenex SecurityGuard cartridge). Mobile phase A was H_2_O/methanol = 95/5 (v/v%), mobile phase B was isopropanol/methanol = 63/37 (v/v%), mobile phase C was H_2_O/methanol/28% NH_4_OH = 93/5/2 (v/v/v%). The LC method consisted of 0.1 mL/min of A/C = 95/5 (v/v%) for 5 min, 0.4 mL/min linear gradient to B/C = 95/5 (v/v%) over 15 min, 0.5 mL/min B/C = 95/5 (v/v%) for 8 min, and equilibration with 0.4 mL/min A/C = 95/5 (v/v%) for 5 min (33 min total run time). The column temperature was 25 °C. For the quantification of BMP, isocratic LC separation was performed with methanol containing 10 mM ammonium formate on a COSMOCORE 2.6C18 column (2.1 × 100 mm; Nacalai Tesque) coupled to an ACQUITY UPLC BEH C18 VanGuard Pre-column (1.7 µm, 2.1 × 5 mm; Waters). The flow rate was 0.3 mL/min and column temperature was 55 °C. For the quantification of other lipid classes, LC separation was performed on an ACQUITY UPLC BEH C18 column (1.7 µm, 2.1 × 100 mm; Waters) coupled to an ACQUITY UPLC BEH C18 VanGuard Pre-column (1.7 µm, 2.1 × 5 mm; Waters). Mobile phase A was acetonitrile/water = 60/40 (v/v%) containing 10 mM ammonium formate and 0.1% (v/v) formic acid, and mobile phase B was isopropanol/acetonitrile = 90/10 (v/v%) containing 10 mM ammonium formate and 0.1% (v/v) formic acid. The LC gradient consisted of 20% B for 2 min, a linear gradient to 60% B over 4 min, a linear gradient to 100% B over 16 min, and equilibration with 20% B for 5 min (total run time: 27 min). The flow rate was 0.3 mL/min, and the column temperature was 55 °C. Multiple reaction monitoring was performed using a Xevo TQ-S micro triple quadrupole mass spectrometry system (Waters) equipped with an ESI source. The ESI capillary voltage was set at 1.0 kV, and the sampling cone was set at 30 V. The source temperature was 150 °C, the desolvation temperature was 500 °C, and the desolvation gas flow was 1,000 L/h. The cone gas flow was 50 L/h.

#### Lipid reaction network analysis

Reaction network analysis was performed as described in the literature.^31^ Using this method, we calculated statistical z scores for lipid reaction pathways to predict whether a particular pathway increased or decreased in *Lamp2b*-Tg compared with WT mice fed HFD for 23 weeks. For constructing a network of reactions, we retrieved reactions that include the lipids identified in our lipidome analyses from the glycerolipid and glycerophospholipid pathways in the KEGG pathway database (https://www.genome.jp/kegg/pathway.html).

#### Measurement of serum lipids and insulin

For serum metabolites analysis, blood was collected from the mice. The samples were then incubated at 25 °C for 1 h, followed by centrifugation (3,000 × g, 20 min) at 4 °C, and serum was collected. The serum T-CHO and TAG levels were measured by Oriental Yeast Co. Ltd. Glucose levels were measured using a glucometer (Accu-Chek Compact; Roche). Serum insulin levels were quantified using an ultra-sensitive mouse insulin ELISA kit (Morinaga Institute of Biological Science, Inc., Cat#49170-54).

#### Glucose and insulin tolerance tests

Prior to the glucose tolerance test, the mice were fasted for 16 h and then injected with glucose (Wako) (1.5 g/kg b.w. i.p. injection). Blood glucose levels were monitored before (0 min) and 15, 30, 60, and 120 min after injection using blood collected from the tail veins and a glucometer (Accu-Chek Compact, Roche). For the insulin sensitivity test, mice were fasted for 4–6 hours and injected with human insulin (Humulin®R Injection) (Eli Lilly Japan) (0.75U/kg b.w. i.p. injection). Tail blood was collected before and at 20, 40, 60, 80, 100, and 120 min after injection, and glucose levels were determined as described above.

### QUANTIFICATION AND STATISTICAL ANALYSIS

Statistical tests of all experiments were performed using GraphPad Prism version Prism 9. For comparisons between two groups, statistical difference was determined by Student’s t test. For comparison of more than two groups, Dunnett’s multiple comparison test or Tukey’s multiple comparisons test was used.

## Notes

### Competing Interest Statement

The authors have declared no competing interest.

