## Supplementary material for "The lysosomal membrane protein LAMP2B mediates microlipophagy to target obesity-related disorders": STAR Methods

**KEY RESOURCES TABLE**

| REAGENT or RESOURCE | SOURCE | IDENTIFIER |
|---|---|---|
| Antibodies | | |
| LAMP2B (1:1000) | outsourced to SCRUM | Cys+LIGRRKTYAGYQTL-OH |
| β-actin (AC-15) (1:10000) | Sigma-Aldrich | Cat#A1978 |
| LC3B (D11) (1:1000) | Cell Signaling | Cat#3868 |
| LAMP2A (1:1000) | Abcam | Cat#ab18528 |
| LAMP2B (1:1000) | Abcam | Cat#ab18529 |
| LAMP2 (H4B4) (1:1000) | Santa Cruz Biotechnology | Cat#sc-18822 |
| LIPA (1:1000) | Novus Biologicals | Cat# NBP1-54155 |
| ATGL (1:1000) | proteintech | Cat# 55190-1-AP |
| α-tubulin | Cell Signaling | Cat#2146 |
| Goat anti-Rabbit IgG (H+L) Secondary Antibody, HRP | Thermofisher | Cat#31460 |
| Goat anti-Mouse IgG (H+L) Secondary Antibody, HRP | Thermofisher | Cat#31430 |
| Chemicals and Peptides | | |
| Insulin-transferrin-sodium selenite media supplement | Sigma | Cat#I3146-5ML |
| dexamethasone | Sigma | Cat#D8893 |
| Streptavidin Sepharose High Performance beads | GE Healthcare | Cat#17-5113-01 |
| TRI Reagent | Sigma-Aldrich | Cat#T9424 |
| Triton-X 100 | Sigma | Cat#T8787-100ML |
| Streptavidin (HRP) | Abcam | ab7403 |
| [$^3$H] oleic acid | PerkinElmer | Product#NET289 |
| oleic acid | nacalai tesque | Cat#25630-51 |
| fatty acid-free BSA | Wako | Cat#013-15143 |
| Triacsin C | Abcam | Cat#ab141888 |
| 4% Paraformaldehyde Phosphate Buffer Solution | Wako | Cat#163-20145 |
| Oil Red O | Sigma | Cat#O0625-25G |
| LysoSensor Green DND-189 | Thermofisher | Cat#L7535 |
| Alexa Fluor-647-Dextran | Thermofisher | Cat# D22914 |
| phosphate buffer (pH 7.4) | LSI Medience Corporation. | Cat#RM102-5L |
| TRI Reagent | MRC | Cat#TR118 |

| | | |
|---|---|---|
| bovine serum albumin | Iwai Chemicals Company | Cat#A001 |
| skim milk | Wako | Cat#190-12865 |
| Chloroquine diphosphate salt | Sigma | Cat#C6628-25G |
| 10% neutral buffered formalin | Wako | Cat#062-01661 |
| Tissue-Tek OCT Compound | Sakura Japan | Cat#4583 |
| D (+) -Glucose | Wako | Cat#041-00595 |
| Humulin R Injection | Eli Lilly Japan | N/A |
| ImmunoStar Zeta | Wako | Cat#295-72404 |
| ImmunoStar LD | Wako | Cat#290-69904 |
| Biotin-conjugated LAMP2A peptides | Fujiwara et al.[17] | N/A |
| Biotin-conjugated LAMP2B peptides | Fujiwara et al.[17] | N/A |
| Biotin-conjugated LAMP2C peptides | Fujiwara et al.[17] | N/A |
| Lipofectamine™ LTX Reagent with PLUS™ Reagent | Thermofisher | Cat#15338100 |
| Lipofectamine RNAiMAX transfection reagent | Thermofisher | Cat#13778150 |
| Critical commercial assays | | |
| QuantiTect Reverse Transcription Kit | Qiagen | Cat#205311 |
| PrimeScript RT Reagent Kit with gDNA Eraser (Perfect Real Time) | TaKaRa | Cat#RR047 |
| SYBR Premix Ex Taq II | TaKaRa | Cat#RR820 |
| Luna® Universal qPCR Master Mix | BioLabs | Cat#M3003 |
| Ultra sensitive Mouse Insulin ELISA kit | Morinaga Institute of Biological Science, Inc. | Cat#49170-54 |
| Deposited data | | |
| Mus musculus, Ensembl 98 | Ensembl database[46] | N/A |
| Glycerolipid metabolism | KEGG[47] | https://www.genome.jp/pathway/map00561 |
| Glycerophospholipid metabolism | KEGG[47] | https://www.genome.jp/pathway/map00564 |
| Experimental models: Cell lines | | |
| HeLa Tet-On cell line | Clontech | Cat#630901 |
| AML12 | ATCC | Cat#CRL-2254 |
| *Atg5*-deficient (*Atg5*$^{-/-}$) mouse embryonic fibroblasts | Kuma et al.[41] | The University of Tokyo, Tokyo, Japan |
| Experimental models: Organisms/strains | | |
| B6C3F1/Slc. | Japan SLC | N/A |
| C57BL/6J Jcl. | CLEA Japan | N/A |

| Oligonucleotides | | |
|---|---|---|
| Human LAMP2A Forward (for q-PCR) | Fujiwara et al.[48] | gggttcagcctttcaatgtg |
| Human LAMP2A Reverse (for q-PCR) | Fujiwara et al.[48] | cagcatgatggtgcttgaga |
| Human LAMP2B Forward (for q-PCR) | Fujiwara et al.[48] | gggttcagcctttcaatgtg |
| Human LAMP2B Reverse (for q-PCR) | Fujiwara et al.[48] | cctgaaagaccagcaccaac |
| Human LAMP2C Forward (for q-PCR) | Fujiwara et al.[48] | gtattctacagctgaagaatgttctg |
| Human LAMP2C Reverse (for q-PCR) | Fujiwara et al.[48] | acacccactgcaacaggaat |
| Human β-actin Forward (for q-PCR) | Takahashi et al.[49] | acaatgtggccgaggacttt |
| Human β-actin Reverse (for q-PCR) | Takahashi et al.[49] | tgtgtggacttgggagagga |
| Mouse Gapdh Forward (for q-PCR) | This paper | tgtgtccgtcgtggatctga |
| Mouse Gapdh Reverse (for q-PCR) | This paper | ttgctgttgaagtcgcaggag |
| Mouse Tnf Forward (for q-PCR) | This paper | ggtgcctatgtctcagcctc |
| Mouse Tnf Reverse (for q-PCR) | This paper | actgatgagagggaggccat |
| Mouse Ccl2; sence | This paper | tcactgaagccagctctctct |
| Mouse Ccl2; antisence | This paper | gtggggcgttaactgcat |
| Mouse Cd68 Forward (for q-PCR) | This paper | ttctgctgtggaaatgcaag |
| Mouse Cd68 Reverse (for q-PCR) | This paper | agaggggctggtaggttgat |
| Mouse Cd163 Forward (for q-PCR) | This paper | acgctggtgtgacatgttct |
| Mouse Cd163 Reverse (for q-PCR) | This paper | cggctacacatcttcccagt |
| Mouse Mrc1 Forward (for q-PCR) | This paper | agaaaatgcacaagagcaagc |
| Mouse Mrc1 Reverse (for q-PCR) | This paper | ggaacatgtgttctgcgttg |
| Mouse Adipoq Forward (for q-PCR) | This paper | caggcatcccaggacatc |
| Mouse Adipoq Reverse (for q-PCR) | This paper | ggaccaagaagacctgcatct |
| siRNA targeting sequence: human LAMP2A; sense | This paper | ggcaggaguacuuauucuagu |
| siRNA targeting sequence: human LAMP2A; antisense | This paper | uagaauaaguacuccugccaa |
| siRNA targeting sequence: human LAMP2B-#1; sense | This paper | gacuaaccccucucuuagagc |
| siRNA targeting sequence: human LAMP2B-#1; antisense | This paper | ucuaagagagggguuagucag |

| | | |
|---|---|---|
| siRNA targeting sequence: human LAMP2B-#2; sense | This paper | cuuaaacaaaaaacuaucaac |
| siRNA targeting sequence: human LAMP2B-#2; antisense | This paper | ugauaguuuuuuguuuaaguu |
| siRNA targeting sequence: mouse Lamp2b-#1; sense | This paper | gcuugauuaucguuauaguga |
| siRNA targeting sequence: mouse Lamp2b-#1; antisense | This paper | acuauaacgauaaucaagccu |
| siRNA targeting sequence: mouse Lamp2b-#2; sense | This paper | gcugauguacguacgauaucu |
| siRNA targeting sequence: mouse Lamp2b-#2; antisense | This paper | auaucguacguacaucagcua |
| siRNA targeting sequence: human LAMP2C; sense | This paper | gguuaucagucuguguaauca |
| siRNA targeting sequence: human LAMP2C; antisense | This paper | auuacacagacugauaaccag |
| siRNA targeting sequence: human LAMP1-#1; sense | This paper | cacacuuucuggcaaacguuu |
| siRNA targeting sequence: human LAMP1-#1; antisense | This paper | acguuugccagaaagugugcc |
| siRNA targeting sequence: human LAMP1-#2; sense | This paper | caguucgggaugaaugcaagu |
| siRNA targeting sequence: human LAMP1-#2; antisense | This paper | uugcauucaucccgaacugga |
| siRNA targeting sequence: human LIPA; sense | This paper | cauacuagcuauuuuuucu |
| siRNA targeting sequence: human LIPA; antisense | This paper | agaaaaaauagcuaguaug |
| siRNA targeting sequence: human ATGL; sense | This paper | guucauugagguaucuaaa |
| siRNA targeting sequence: human ATGL; antisense | This paper | uuuagauaccucaaugaac |
| siRNA targeting sequence: mouse Vps4a; sense | This paper | caacccuagcguuaugauu |
| siRNA targeting sequence: mouse Vps4a; antisense | This paper | aaucauaacgcuaggguug |
| siRNA targeting sequence: mouse Vps4b; sense | This paper | cuaccuugcgagugcuaca |
| siRNA targeting sequence: mouse Vps4b; antisense | This paper | uguagcacucgcaagguag |
| siRNA targeting sequence: mouse Tsg101; sense | This paper | gguacaaucccagugcguu |
| siRNA targeting sequence: mouse Tsg101; antisense | This paper | aacgcacugggauuguacc |
| siRNA targeting sequence: mouse Alix; sense | This paper | ggauuacuuuggcgaugcu |
| siRNA targeting sequence: mouse Alix; antisense | This paper | agcaucgccaaaguaaucc |
| siRNA targeting sequence: mouse Chmp4b; sense | This paper | gaaacagucccucuaccaa |

| | | |
|---|---|---|
| siRNA targeting sequence: mouse Chmp4b; antisense | This paper | uugguagagggacuguuuc |
| siRNA targeting sequence: enhanced green fluorescent protein (EGFP)-#1; sense | Aizawa et al.[50] | gccacaacgucuauaucaugg |
| siRNA targeting sequence: enhanced green fluorescent protein (EGFP)-#1; antisense | Aizawa et al.[50] | augauauagacguuguggcug |
| siRNA targeting sequence: enhanced green fluorescent protein (EGFP)-#2; sense | Aizawa et al.[50] | cagcacgacuucuucaagucc |
| siRNA targeting sequence: enhanced green fluorescent protein (EGFP)-#2; antisense | Aizawa et al.[50] | acuugaagaagucgugcugcu |
| siRNA targeting sequence: universal negative control (UNC); sense | This paper | uucuccgaacgugucacgu |
| siRNA targeting sequence: universal negative control (UNC); antisense | This paper | acgugacacguucggagaa |
| Plasmid DNA | | |
| pCI-neo-human LAMP2A | Kabuta et al.[51] | N/A |
| pCI-neo-human LAMP2B | Fujiwara et al.[17] | N/A |
| pCI-neo-human LAMP2B-RRSS | This paper | N/A |
| pCI-neo-human LAMP2B-L410A | This paper | N/A |
| pCI-neo-human LAMP2C | This paper | N/A |
| pEGFP-N1- human LAMP2B-GFP | This paper | N/A |
| pEGFP-N1- human LAMP2B-RRSS-GFP | This paper | N/A |
| pEGFP-N1- human LAMP2B-L410A-GFP | This paper | N/A |
| pTag-BFP-N-human LAMP1-BFP | This paper | N/A |
| pmCherry-C1-EGFP-mPLIN2 | This paper | N/A |
| Software and algorithms | | |
| ImageJ | NIH | https://imagej.net/ |
| FactoMineR | Le et al.[52] | https://cran.r-project.org/web/packages/FactoMineR/index.html |
| factoextra | https://rpkgs.datanovia.com/factoextra/index.html | https://cran.r-project.org/web/packages/factoextra/index.html |
| iheatmapr | Schep and Kummerfeld[53] | https://cran.r-project.org/web/packages/iheatmapr/index.html |
| GraphPad Prism 9 | GraphPad | https://www.graphpad.com/features |

| Instrumentation | | |
|---|---|---|
| Accu-Chek Compact Plus | Roche | Cat#000102 |
